# *Staphylococcus aureus* uses a eukaryotic-like uridyltransferase to make UDP-GlcNAc for cell wall synthesis

**DOI:** 10.64898/2026.06.23.733835

**Authors:** Pak-Ming Fong, Anna I. Weaver, Abigail L. Manson, Youngseon Park, Ashlee M. Earl, Suzanne Walker

## Abstract

UDP-N-acetylglucosamine (UDP-GlcNAc) is an essential metabolite used to build the peptidoglycan cell wall that protects bacteria against lysis caused by high turgor pressure. More than thirty years ago, GlmU, which contains both acetyltransferase and uridyltransferase activities, was discovered to make UDP-GlcNAc in *E. coli,* and it is now widely considered essential for UDP-GlcNAc synthesis across bacteria. Here we report that the clinically important pathogen *Staphylococcus aureus* relies on a previously uncharacterized uridyltransferase called NagU to synthesize most of its UDP-GlcNAc for cell wall synthesis. We also report that NagU binds to and negatively regulates DacA, the diadenylate cyclase that makes the essential second messenger cyclic di-AMP, which regulates turgor pressure. We propose a model in which increasing NagU expression simultaneously increases flux into peptidoglycan and increases turgor pressure to enable cell wall expansion. Finally, an evolutionary analysis shows that many bacteria beyond Staphylococcaceae encode NagU homologues but lack GlmU either entirely or in part, suggesting that alternative UDP-GlcNAc biosynthesis pathways are not rare.

## INTRODUCTION

Bacterial cells are surrounded by a multi-layered cell envelope that enables them to survive in diverse and often hostile environments. The universally conserved component of the bacterial cell envelope is the peptidoglycan cell wall, a mesh-like structure that surrounds the cytoplasmic membrane, providing mechanical strength that counteracts high turgor pressure and prevents membrane rupture^1, 2^. Bacterial viability depends on maintaining the integrity of the cell wall^3, 4^. Bacteria and fungi have evolved to exploit this vulnerability to gain a competitive advantage, producing a diverse range of natural products that target different steps in peptidoglycan assembly^5–7^. Because peptidoglycan biosynthesis has been extensively validated as an antibiotic target, virtually every known cell wall enzyme in well-studied pathogens has been scrutinized as a new possible target^8, 9^. Given this context, the discovery of a new peptidoglycan biosynthesis enzyme in an intensively studied pathogen such as *Staphylococcus aureus* is unexpected. Nevertheless, we report here that *S. aureus* and other members of the family Staphylococcaceae harbor a previously overlooked enzyme that makes a precursor essential to build cell wall peptidoglycan.

Across bacteria, peptidoglycan is composed of glycan chains made of repeating disaccharide units consisting of *N*-acetylglucosamine (GlcNAc) and *N*-acetylmuramic acid (MurNAc). These chains are held together by cross-bridges between peptides attached to the MurNAc moieties^1, 2^. Both of the sugars in the repeating disaccharide subunit of peptidoglycan are derived from a single precursor, UDP-GlcNAc. Canonically, the bifunctional enzyme GlmU is responsible for making UDP-GlcNAc from glucosamine-1-phosphate. GlmU’s C-terminal domain first acetylates the C2 amine of glucosamine-1-phophate and then its N-terminal domain uridylates the anomeric phosphate^10, 11^. GlmU was discovered in *E. coli* in the early 1990s where it was found to be essential^12, 13^. It is broadly conserved in other bacteria, including *S. aureus* where the *glmU* gene is also essential. Although it has been assumed that both the acetylation and uridylation activities of GlmU are required in *S. aureus*, this assumption has never been tested.

*S. aureus* encodes a predicted monofunctional uridyltransferase (SAOUHSC_02423), here named NagU (for <u>N</u>-<u>a</u>cetyl<u>g</u>lucosamine <u>u</u>ridyltransferase), that has not been previously characterized. Curious about NagU’s role, we investigated it and have found that it makes UDP-GlcNAc both *in vitro* and in cells. Surprisingly, our data show that most of the UDP-GlcNAc *S. aureus* makes is due to NagU, not to GlmU’s uridyltransferase activity. NagU thus becomes essential for viability under cell envelope stress. We have also found that NagU directly interacts with and negatively regulates DacA, which synthesizes the important secondary messenger cyclid-di-AMP (c-di-AMP)^14^. Since c-di-AMP regulates turgor pressure, with lower levels resulting in higher turgor^15, 16^, we propose that NagU coordinates turgor pressure with the tensile strength provided by peptidoglycan. We suggest that higher turgor is required under conditions that demand higher precursor flux into peptidoglycan because turgor-promoted cell wall expansion makes space to incorporate nascent glycan chains into the existing cell wall matrix.

## RESULTS

### NagU is a eukaryotic-like uridyltransferase that specifically makes UDP-GlcNAc *in vitro*

UDP-GlcNAc is essential in both prokaryotes and eukaryotes but is canonically made by different enzymatic pathways in these kingdoms. In bacteria, UDP-GlcNAc is produced via a conserved three-enzyme pathway comprising four sequential reactions. GlmS synthesizes glucosamine-6-phosphate (GlcN-6-P) from fructose-6-phosphate (F-6-P) and glutamine, and GlmM isomerizes GlcN-6-P to glucosamine-1-phosphate (GlcN-1-P). The bifunctional enzyme GlmU then catalyses the final two reactions: acetylation of the C2 amine of GlcN-1-P to produce *N*-acetylglucosamine-1-phosphate (GlcNAc-1-P) followed by uridylation of GlcNAc-1-P to produce UDP-GlcNAc (Fig. 1a)^17^. In eukaryotes, glucosamine-6-phosphate is acetylated on the C2 amine by a monofunctional acetyltransferase before the phosphate moves to the anomeric carbon, and the final step in the pathway, uridylation of GlcNAc-1-P, is catalyzed by the monofunctional uridyltransferase UAP1 (also known as AGX1) (Fig. 1a)^18–20^. Like most bacteria *S. aureus* contains GlmU. However, this organism also contains a previously unstudied gene, *nagU*, which was annotated as a uridyltransferase. We used a Foldseek search against the PDB database to look for proteins that are structurally similar to the predicted structure of the encoded protein^21^ (Supplementary Table 2). The top hit was the human UAP1, which synthesizes UDP-GlcNAc from GlcNAc-1-P (Fig. 1b).

**Fig. 1.**
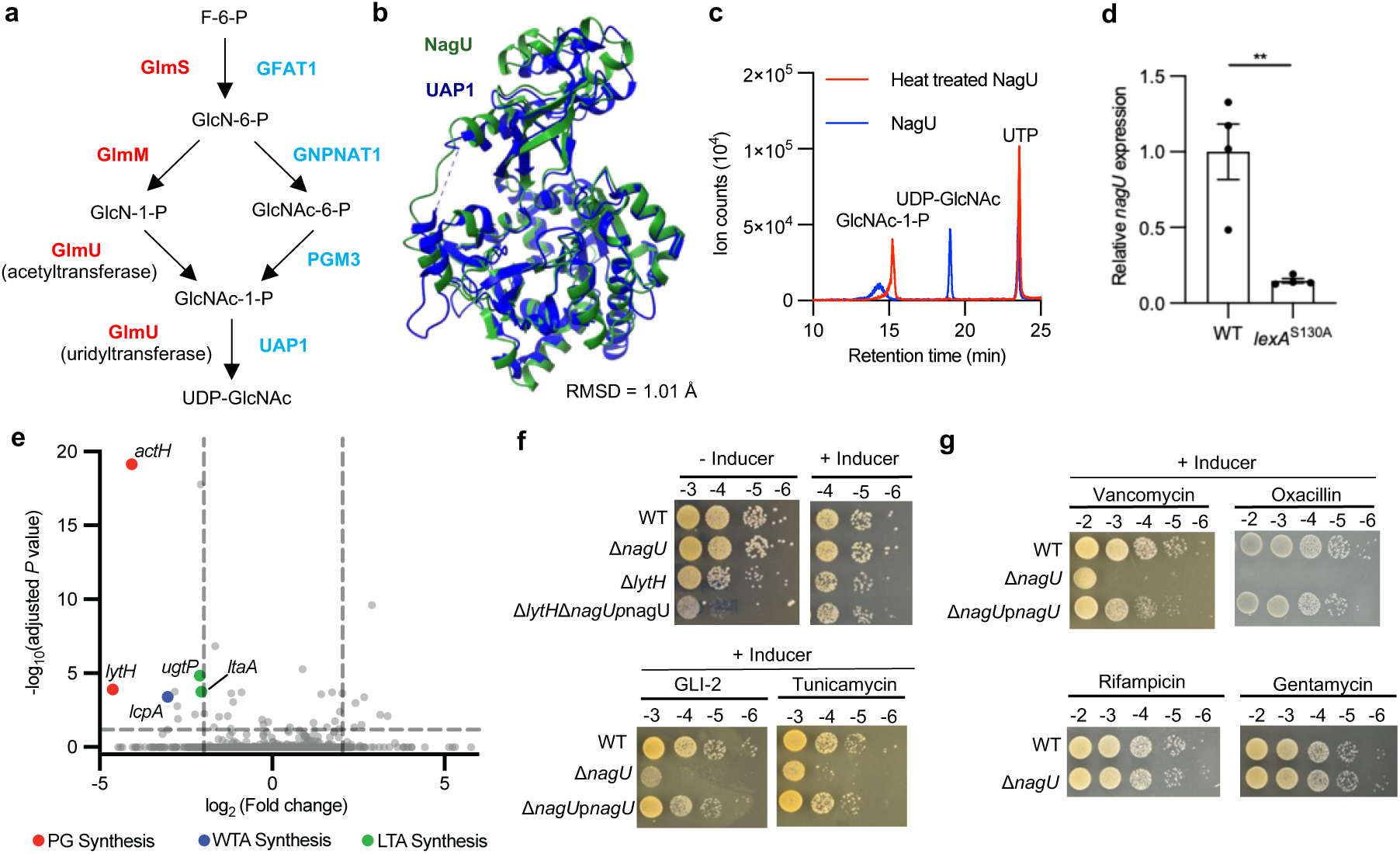
NagU is a uridyltransferase to synthesize UDP-GlcNAc required for survival under cell envelope stress. **a,** Schematic of the UDP-GlcNAc biosynthetic pathway in prokaryotes (red) and eukaryotes (blue). **b**, Structural alignment of the AlphaFold predicted structure of NagU (AF-Q2FW81-F1) and the crystal structure of human UAP1 (PDB: 1JV3_B). **c**, Liquid chromatography mass spectrometry (LCMS) traces of uridyltransfer reaction using native or heat-treated NagU with *N*-acetylglucosamine-1-phosphate (GlcNAc-1-P) and UTP as substrates. **d**, RT-qPCR analysis of *nagU* expression relative to internal reference *rpoB* in WT HG003 and *lexA*^S130A^ strains during early log phase. The average of the relative expression in WT is set to 1.0. Data are means ± SEM of 4 biological replicates. \*\**P* < 0.01, ns,. Statistical significance was determined by two-tailed unpaired Student’s *t* test. **e**, Volcano plot of transposon insertions within individual genes in a Δ*nagU* transposon library relative to a WT control library. Significantly depleted genes (>4-fold, adjusted P < 0.05) related to peptidoglycan (PG), wall teichoic acid (WTA), and lipoteichoic acid (LTA) synthesis are labelled. Statistical significance was determined by two-sided Mann-Whitney U test corrected for multiple hypothesis testing via the Benjamini-Hochberg method. **f**. Spot titer assays evaluating the genetic interaction between *nagU* and *lytH* (Top; inducer = 0.4 µM anhydrotetracycline), and assessing sensitivity of Δ*nagU* to the UgtP inhibitor GLI-2 (2.0 µg mL⁻¹; Bottom left) and the TarO inhibitor tunicamycin (0.2 µg mL⁻¹; Bottom right). Strains spotted include WT HG003, Δ*nagU*, Δ*lytH*, Δ*nagU*Δ*lytH* with *nagU*-complemented strain (Top) and WT HG003, Δ*nagU*, Δ*nagU* with *nagU*-complemented strain (Bottom). **g,** Spot titer assays evaluating the sensitivity of Δ*nagU* to cell wall-targeting antibiotics. Strains spotted include WT HG003, Δ*nagU*, Δ*nagU* Δ*nagU* with *nagU*-complemented strain. Strains were spotted on TSA containing 0.725 µg mL⁻¹ vancomycin (Top left), 0.1 µg mL⁻¹ oxacillin (Top right), 1 ng mL⁻¹ rifampicin (Bottom left) or 1.25 µg mL⁻¹ gentamycin (Bottom right). Inducer = 0.4 µM anhydrotetracycline.

We tested if *S. aureus* NagU has uridyltransferase activity *in vitro*. To do so, we purified NagU and assessed product formation using liquid chromatography mass spectrometry (LCMS). NagU, but not the heat-inactivated enzyme, converted GlcNAc-1-P and UTP into UDP-GlcNAc (Fig. 1c). We also tested glucose-1-phosphate (Glc-1-P) as a potential substrate because *S. aureus* uses UDP-Glc to make another important cell envelope polymer, lipoteichoic acid (LTA)^22^. However, NagU was unable to convert this substrate to the corresponding activated sugar (Extended Data Fig. 1). We concluded that NagU is a uridyltransferase with selectivity for converting GlcNAc-1-P to UDP-GlcNAc.

### *nagU* is abundantly expressed under normal growth conditions in *S. aureus*

NagU is encoded in an operon that has a canonical SOS box in the promoter region, suggesting that it is required for the SOS response^23, 24^. To examine whether *nagU* is under the control of the transcriptional repressor LexA, which represses SOS responsive genes, we used real-time quantitative PCR (RT-qPCR) to compare *nagU* expression in wildtype *S. aureus* to expression in a *lexA*^S130A^ mutant, which expresses a LexA variant that cannot be cleaved to derepress gene expression^23, 25^. We found that *nagU* expression was lower in the *lexA*^S130A^ mutant compared to wildtype (Fig. 1d). We also found that *nagU* expression increased under ciprofloxacin treatment in the wild-type strain compared to the *lexA*^S130A^ mutant (Extended Data Fig. 2a). These experiments showed that LexA regulates the expression of *nagU*, as predicted. However, the Δ*nagU* strain was not hypersensitive to ciprofloxacin compared with wildtype *S. aureus* (Extended Data Fig. 2b). Moreover, there appears to be substantial basal expression of *nagU* even in the absence of DNA damage induction (Fig. 1d). Consistent with this, ribosome profiling data shows that NagU expression levels are comparable to those of many other metabolic enzymes.^26^ Given these observations, we tentatively concluded that NagU plays a role in *S. aureus* physiology that is independent of DNA damage signalling.

### A TnSeq screen implicates NagU in cell envelope physiology

TnSeq screens can provide useful information about general functions of poorly characterized genes. We therefore made a transposon mutant library in a Δ*nagU* background and compared the transposon insertion reads across the genome to those for the corresponding wildtype library. The hits in the screen were mainly found in cell envelope biosynthesis pathways (Fig. 1e, Extended Data Fig. 3a, Supplementary Table 1). For example, the top depleted hits were *lytH* and *actH*, which encode a membrane-anchored amidase complex important in the peptidoglycan biosynthesis pathway^27, 28^. Other depleted hits included *lcpA*, which encodes the principal ligase that attaches wall teichoic acid (WTA) to peptidoglycan^29, 30^, and two genes involved in LTA anchor assembly, *ugtP* and *ltaA*^22, 31–33^. We examined synthetic phenotypes between Δ *nagU* and selected pathways identified in the screen using genetic and pharmacological methods. We found that depleting *nagU* in a Δ*lytH* background resulted in a synthetic sick phenotype, as did plating Δ*nagU* cells on the UgtP inhibitor GLI-2^34^ or the WTA inhibitor tunicamycin^35^ (Fig. 1f, Extended Data Fig. 3b). Taken together, the hits suggested an important role for NagU in cell envelope integrity, a role that would be consistent with its enzymatic function in making UDP-GlcNAc.

### Cells lacking NagU are sensitive to antibiotics that target the cell wall

Because the TnSeq screen implicated NagU in cell envelope integrity, we examined the sensitivity of Δ*nagU* to a range of antibiotics having different targets. In spot titer assays, Δ*nagU* behaved similarly to wildtype *S. aureus* on rifampicin, gentamycin, and ciprofloxacin, which do not target the cell wall (Fig. 1g, Extended Data Fig. 3c); however, it was more sensitive than wildtype on vancomycin, oxacillin, daptomycin, and moenomycin, all of which target peptidoglycan assembly (Fig. 1g and Extended Data Fig. 3c and d). Because UDP-GlcNAc is needed to make bacterial peptidoglycan, we also examined whether deleting or overexpressing *nagU* affected cell wall thickness. Transmission electron micrographs of cells lacking or overexpressing *nagU* showed that cell wall thickness at the cell periphery was thinner in Δ*nagU* cells and thicker in cells overexpressing *nagU* than in WT cells (Extended Data Fig. 4a and b). These results are consistent with a role for *nagU* in making UDP-GlcNAc to build the cell wall.

### NagU and GlmU have redundant enzymatic functions in cells

Because *S. aureus* harbours a eukaryotic-like uridyltransferase that makes UDP-GlcNAc and is functionally important for cell envelope integrity, we asked if GlmU’s uridyltransferase activity is essential in this organism. To address this question, we constructed *glmU*^R15A^, which contains a mutation in the native locus of *glmU*. A previous study showed that mutating the corresponding arginine in *Escherichia coli* GlmU resulted in a more than 6000-fold reduction in enzymatic activity and a loss of viability^36^. In *S. aureus*, the *glmU*^R15A^ mutant was fully viable, suggesting that another enzyme compensates for the loss of GlmU’s uridyltransferase function (Fig. 2a). To assess if *nagU* is essential in the *glmU*^R15A^ mutant, we introduced an inducible copy of *nagU* at an ectopic locus and then deleted the native copy of *nagU*. We found that this strain was unable to grow in the absence of inducer (Fig. 2a), showing that NagU is essential if GlmU’s uridylation domain is inactivated. Therefore, NagU functions as a GlcNAc-1-P uridyltransferase *in vivo* as well as *in vitro*. We note that a previous study identified *S. aureus* GlmR (encoded by *saouhsc_00788*; also known as *yvck*) as a GlcNAc-1-P uridyltransferase *in vitro*, but it has not been studied *in vivo*^37^. However, our experiments show that natively regulated GlmR cannot compensate for the combined loss of the uridyltransferase activities of NagU and GlmU; it therefore remains unclear if GlmR has uridyltransferase activity in a cellular context.

**Fig. 2.**
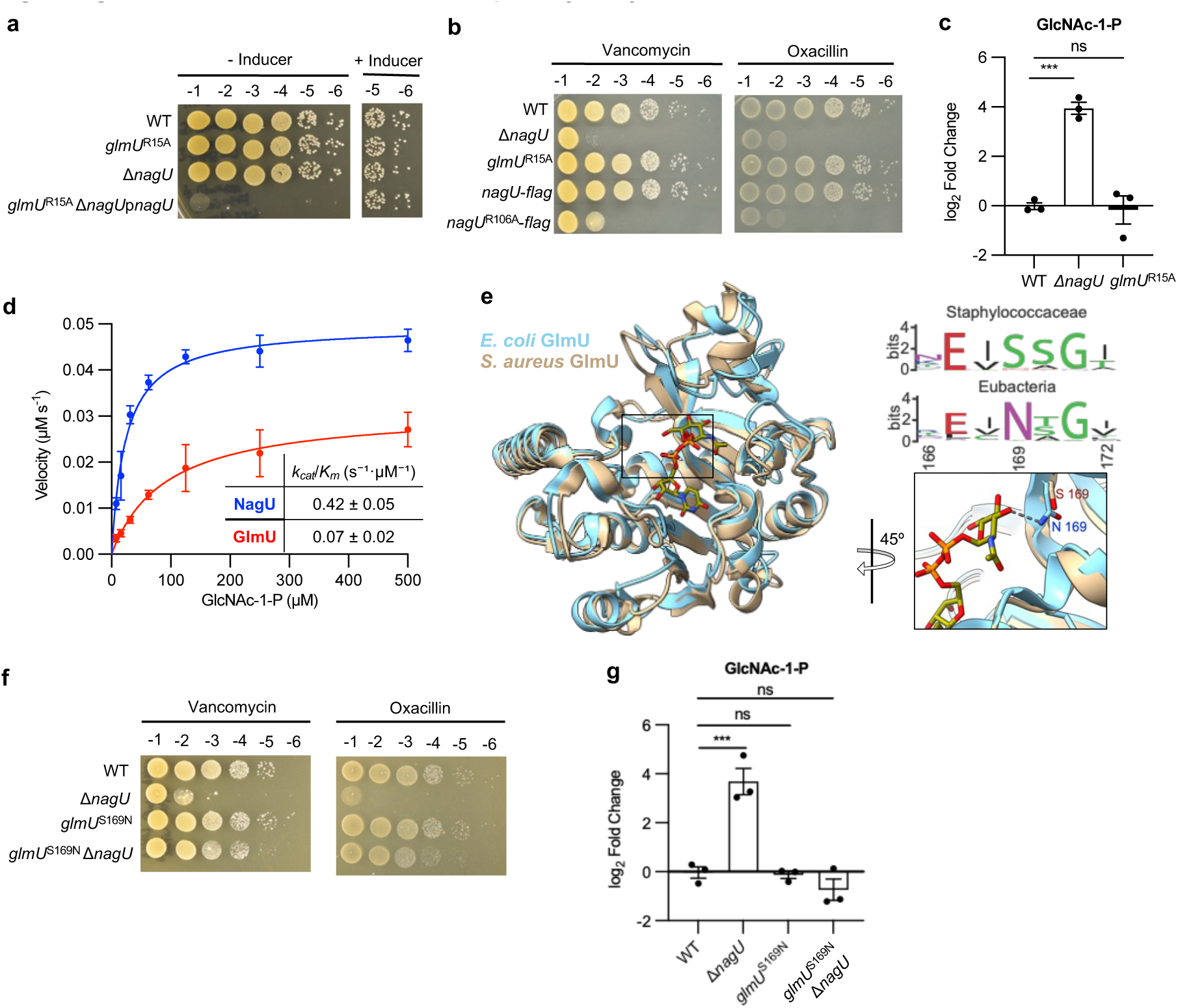
NagU, rather than GlmU, serves as the primary uridyltransferase in *S. aureus*. **a**, Spot titers of *S. aureus* strains harbouring an inactive GlmU uridyltransferase in WT and Δ*nagU* backgrounds. Strains spotted include WT HG003, strains with chromosomally mutated *glmU* encoding inactive uridyltransferase (*glmU*^R15A^), Δ*nagU* and Δ*nagUglmU*^R15A^ with *nagU*-complemented strain. Inducer = 0.4 µM anhydrotetracycline. **b**, Spot titer assay evaluating oxacillin and vancomycin sensitivity following the inactivation of uridylation activity of NagU or GlmU. Strains include WT, Δ*nagU*, *glmU*^R15A^, *nagU*-*flag* encoding NagU with C-terminal FLAG tag (LF1287), and *nagU*^R106A^ encoding inactive uridyltransferase with C-terminal FLAG tag (LF1289). Strains were spotted on TSA containing 0.725 µg mL⁻¹ vancomycin (Left), 0.1 µg mL⁻¹ oxacillin (Right). **c**, UPLC quantification of cellular GlcNAc-1-P levels. Data are means ± SEM of 3 biological replicates. \*\*\**P* < 0.001, ns, not significant; Statistical significance was determined by one-way ANOVA with Dunnett corrections. Strains include WT, Δ*nagU* and *glmU*^R15A^. **d**, Michaelis-Menten kinetics of NagU-FLAG and His_6_-GlmU using GlcNAc-1-P as substrate. *k_cat_*/*K_m_* are means ± SEM of 3 biological replicates (See Supplementary table 3). **e**, Structural alignment of the crystal structures of *E. coli G*lmU (Blue, PDB: 1hv9) and *S. aureus* GlmU (Brown, PDB: 9DQF) (Left) highlighting the catalytic site (Bottom right boxed region), alongside a Sequence Logo of GlmU homologs in Staphylococcaceae and in all Eubacteria (Top right). **f**, Spot titer assay evaluating the viability of the *glmU*^S169N^ mutation in WT and Δ*nagU* under antibiotic stress. Strains include WT HG003, Δ*nagU*, *glmU*^S169N^, and *glmU*^S169N^Δ*nagU*. Strains were spotted on TSA containing 0.1 µg mL⁻¹ oxacillin and 0.725 µg mL⁻¹ vancomycin. **g**. UPLC quantification of cellular GlcNAc-1-P levels for the strains described in **f**. Data are means ± SEM of 3 biological replicates. \*\*\**P* < 0.001, ns, not significant; Statistical significance was determined by one-way ANOVA with Dunnett’s correction.

### NagU is the predominant uridyltransferase in *S. aureus*

The results above showed that *S. aureus* harbours two redundant uridyltransferases for making UDP-GlcNAc, but whether one is more active – and therefore more important for cell wall synthesis - was unclear. To assess the relative importance of these distinct uridyltransferases, we compared phenotypes of cells expressing either *nagU*^WT^*glmU*^R15A^ or *nagU*^R106A^*glmU*^WT^. The *nagU* mutation was based on studies of the NagU yeast ortholog, UAP1, which showed that R106 is required for activity^20^. To confirm NagU inactivation, we purified NagU^R106A^ and tested its uridylation activity in vitro. Consistent with the prediction, the uridyltransferase activity of NagU^R106A^ was undetectable (Extended Data Fig. 5a). We constructed *nagU*^R106A^ at the native locus like the *glmU*^R15A^ mutant so that both genes are under native expression. After verifying that protein stability was unaffected by the mutations (Extended Data Fig. 5b), we plated strains on oxacillin and vancomycin, two conditions to which Δ*nagU* is hypersensitive. The spot assays showed that *nagU*^R106A^*glmU*^WT^ was hypersensitive to oxacillin and vancomycin just like Δ*nagU*. However, the viability of the *nagU*^WT^*glmU*^R15A^ strain was comparable to wildtype under these conditions (Fig. 2b). Hence, under cell wall stress, NagU’s uridyltransferase activity is more important than GlmU’s. We interpreted these findings to mean that NagU is responsible for more UDP-GlcNAc synthesis in cells than GlmU.

We used an orthogonal approach to more directly assess the uridyltransferase activity of NagU and GlmU in cells. We extracted polar metabolites from *S. aureus* WT and mutants lacking either NagU or GlmU’s uridylation activity and used ultra-performance liquid chromatography-mass spectrometry (UPLC-MS) to quantify the steady-state levels of metabolites in the biosynthesis pathways for cell wall precursors. These metabolomics assays are demanding and processing steps can obscure small differences in steady-state metabolite levels between strains. Perhaps for this reason, we did not observe a statistically significance difference in levels for most metabolites. However, GlcNAc-1-P accumulated more than 8-fold in the Δ*nagU* and *nagU*^R106A^ strains compared to wild type, but was unchanged in the *glmU*^R15A^ strain (Fig. 2c, Extended Data Fig. 5c). The large accumulation of GlcNAc-1-P in the *nagU* mutants is consistent with a bottleneck in flux to UDP-GlcNAc that does not occur when the uridyltransferase activity of GlmU is lost. Taken together with the antibiotic susceptibility profiles, these results provide compelling evidence that NagU is the primary uridyltransferase that makes UDP-GlcNAc in *S. aureus*.

### *S. aureus GlmU* has an impaired active site

NagU is the primary uridyltransferase in *S. aureus*, but we wondered if it has higher intrinsic activity than GlmU or is simply expressed at higher levels. Published ribosome profiling data suggests that NagU is expressed at modestly higher levels than GlmU^26^, and qRT-PCR data is consistent with this (Extended Data Fig. 6a). However, the expression differences did not seem large enough to explain the substantial accumulation of GlcNAc-1-P in *nagU* mutants. Therefore, we assessed the biochemical activities of NagU and GlmU *in vitro* by monitoring pyrophosphate formation during the uridyltransfer reaction. Steady-state kinetic parameters from a Michaelis-Menten analysis of GlcNAc-1-P turnover showed that the catalytic rate constant for NagU is about 1.6-fold faster than for GlmU (9.9 s^−1^ compared with 6.3 s^−1^), while *k_cat_*/*K_m_* is approximately 6-fold higher (0.42 s⁻¹·µM⁻¹ compared with 0.07 s⁻¹·µM⁻¹) (Fig. 2d, Supplementary Table 3). A similar trend is observed for UTP (Extended Data Fig. 6b, Supplementary Table 3). We concluded that the dominance of NagU *in vivo* is driven largely by its superior intrinsic enzymatic activity (higher *k_cat_* and lower *Km*).

The activity differences between the two enzymes were large enough that we wondered if the uridyltransferase activity of GlmU was defective in *S. aureus.* Therefore, we compared GlmU sequences across bacteria to determine if there were notable differences. Although most of the highly conserved GlmU residues are also conserved in *S. aureus* GlmU, the *S. aureus* enzyme contains a serine rather than an asparagine in a conserved active site motif (Ser169). Almost all other members of the family Staphylococcaceae similarly have a serine at this position (Fig. 2e, Supplementary Table 4). A crystal structure of *E. coli* GlmU with UDP-GlcNAc bound shows that the conserved active site asparagine makes an apparent H-bond to the C3’-OH position of the UDP-GlcNAc (Fig. 2e)^10, 36^. We therefore hypothesized that the serine substitution at position 169 of *S. aureus* GlmU was largely responsible for its reduced uridyltransferase activity. To examine if replacing the serine with asparagine would increase GlmU’s activity, we mutated *glmU*^WT^ to *glmU*^S169N^ in both wildtype and Δ*nagU* backgrounds and plated the strains on oxacillin and vancomycin. We found that *glmU*^S169N^ rescued growth of Δ*nagU* on both oxacillin and vancomycin (Fig. 2f), indicating that the amino acid change had increased GlmU’s uridylation activity. Consistent with this, targeted metabolomics showed that GlcNAc-1-P did not accumulate in the Δ*nagU glmU*^S169N^ strain like it did in the Δ*nagU strain* (Fig. 2g). Due to aggregation, we were unable to purify GlmU^S169N^ in a form that would enable us to compare its enzymatic activity to GlmU WT. Nevertheless, the ability of this single substitution to rescue the growth of Δ*nagU* under cell envelope stress and prevent GlcNAc-1-P accumulation strongly supports the conclusion that the GlmU uridyltransferase activity in *S. aureus* and other Staphylococci is compromised by this deleterious active site mutation. Accordingly, these organisms depend primarily on NagU to make sufficient UDP-GlcNAc to meet demands when cells are under envelope stress.

### NagU physically interacts with the diadenylate synthase DacA

We asked if NagU has other functions that might help explain why *S. aureus* and related bacteria have come to rely on it for UDP-GlcNAc synthesis. To determine if NagU has a protein partner that would reveal another possible function, we overexpressed NagU with a C-terminal FLAG tag in Δ*nagU* cells and performed a immunoprecipitation assay to capture protein interactors. In repeated immunoprecipitations, we identified the diadenylate synthase DacA and its regulatory protein YbbR (Fig. 3a, Extended Data Fig. 7a, Supplementary Table 5). DacA is an essential enzyme that makes the bacterial second messenger cyclic-di-AMP (c-di-AMP). In Gram-positive organisms, c-di-AMP controls turgor pressure and both high and low levels can be deleterious, so DacA is tightly regulated via multiple mechanisms^37–41^. We used *in silico* modelling to assess how NagU might bind to DacA or YbbR. AlphaFold predicted that the N-terminal domain of NagU binds directly to the cytosolic domain DacA whereas no interaction was predicted with YbbR (Fig. 3b, Extended Data Fig. 7b and 7c). To confirm the interaction, we separately purified NagU-FLAG and the cytosolic domain of DacA (His_6_-DacA_cd_)^42^ and incubated them prior to analysis by size exclusion chromatography coupled with multi-angle light scattering (SEC-MALS) (Fig. 3c, Extended Data Fig. 7d). We observed a 1:1 and a 1:2 NagU:DacA complex, but not a 2:2 complex even though one would be expected based on symmetry and AlphaFold predictions (Extended Data Fig. 7c). The binding affinity may be insufficient for the 2:2 complex to survive the SEC-MALS analysis in intact form. Nevertheless, these studies confirmed that NagU interacts with DacA, suggesting it might modulate cellular c-di-AMP production.

**Fig. 3.**
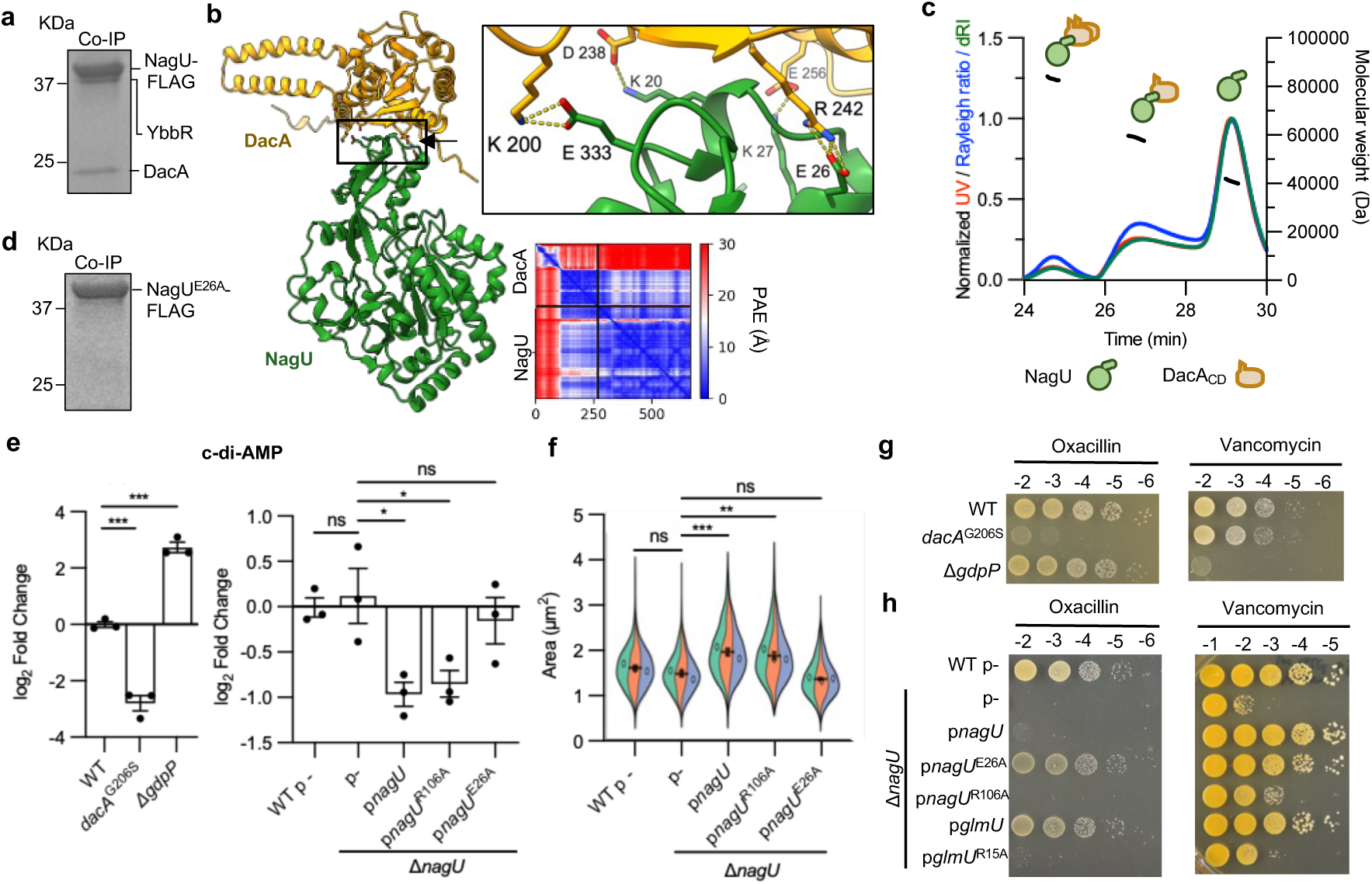
NagU physically interacts with the c-di-AMP synthase DacA to negatively regulate c-di-AMP synthesis. **a**, Coomassie-stained gel of eluate from a immunoprecipitation using NagU-FLAG as bait. The bands were subjected to LC-MS to identify NagU’s protein partners. **b**, AlphaFold2-predicted structure of NagU-DacA complex and predicted polar contacts mediating the interaction (Top right boxed region). The position alignment error (PAE) plot is shown below (Bottom right). **c**, SEC-MALs chromatogram of NagU and DacA_CD_ showing complexes and NagU monomer. **d**, Coomassie-stained gel of eluate from a immunoprecipitation using NagU^E26A^ mutant as bait. **e**, UPLC quantification of cellular c-di-AMP in the hypomorphic strain *dacA*^G206S^ and Δ*gdpP* (Left), and in strains overexpressing WT *nagU* and variants (Right). Strains include WT, *dacA*^G206S^ and Δ*gdpP* (Left), WT carrying an empty vector, and Δ*nagU* carrying an empty vector or vectors expressing *nagU*, *nagU*^R106A^ and *nagU*^E26A^ (Right). 1 mM IPTG was added to induce overexpression. Data are means ± SEM of 3 biological replicates \*\*\**P* < 0.001, \**P* < 0.05, ns, not significant; Statistical significance was determined by one-way ANOVA with Dunnett’s correction. **f**, Cell size quantification of the strains wildtype carrying empty vector, Δ*nagU* carrying empty vector or vector expressing *nagU*, *nagU*^R106A^ and *nagU*^E26A^. Inducer 1 mM IPTG was added for overexpression. Cells were stained with the membrane dye Nile Red, and volumes of nondividing cells were quantified and plotted as Violin SuperPlots. \*\*\**P* < 0.001, \*\**P* < 0.01, ns, not significant; Statistical significance was determined by a one-way ANOVA with Bonferroni corrections. **g**, Spot titer evaluating the sensitivity of *dacA*^G206S^ and Δ*gdpP* to oxacillin (Left) and vancomycin (Right). Strains were spotted on TSA plates containing 0.1 µg mL⁻¹ oxacillin or 0.325 µg mL^−1^ vancomycin. **h**, Spot titer assays evaluating the antibiotic sensitivity of strains overexpressing *nagU*, *glmU* and their variants to oxacillin and vancomycin. Strains spotted include WT carrying empty vector, Δ*nagU* carrying empty vector or vector expressing *nagU*, *nagU*^R106A^*, nagU*^E26A^, *glmU* and *glmU*^R15A^. Strains were spotted on TSA plates containing 10 µg mL⁻¹ erythromycin and 0.1 µg mL⁻¹ oxacillin or 0.325 µg mL⁻¹ vancomycin. 1 mM IPTG was added for overexpression.

### NagU reduces cellular c-di-AMP via interaction with the diadenylate synthase DacA

We wondered if NagU can modulate c-di-AMP production in cells. To address this question, we required a catalytically active NagU variant incapable of binding to DacA. In the AlphaFold model, four NagU residues form polar contacts to DacA (Fig. 3b). Of these, Glu26 is centrally located at the binding interface and is predicted to form a bidentate H bond with Arg 242 in DacA. Because removing Glu26 would result in the loss of two predicted H-bonds, we prioritized changing this residue to destabilize the complex. NagU^E26A^ had full catalytic activity *in vitro*, which was not surprising given that the mutation is in a separate domain from the active site (Extended Data Fig. 5a). Moreover, NagU^E26A^ expressed similarly to wild type (Extended Data Fig. 5b). Steady-state levels of GlcNAc-1-P were also the same as wild type in the *nagU*^E26^ variant, showing that it has full catalytic activity in cells (Extended Data Fig. 5c). However, unlike wild type NagU, NagU^E26A^ was unable to immunoprecipitate DacA and YbbR (Extended Data Fig. 8a, Supplementary Table 5). In contrast, NagU^R106A^, which has an inactivating mutation in the active site, behaved just like wildtype NagU with respect to its ability to immunoprecipitate DacA/YbbR (Extended Data Fig. 8a, Supplementary Table 5). We concluded from these experiments that AlphaFold accurately predicted the NagU-DacA interface and that we had successful engineered a catalytically active NagU variant that is unable to form a complex with DacA.

We next assessed whether NagU binding to DacA influences c-di-AMP production. Using the UPLC-MS assay, we compared c-di-AMP levels in strains overexpressing *nagU*^E26A^ to those for NagU WT and the catalytically dead *nagU*^R106A^ variant. As control strains, we used the *dacA* hypomorph *dacA*^G206S^ and a mutant lacking *gdpP*, the phosphodiesterase that degrades c-di-AMP^40, 43^. As expected, the hypomorphic *dacA*^G206S^ strain had low cellular c-di-AMP compared to wildtype while the Δ*gdpP* strain had elevated c-di-AMP (Fig. 3e). There was no significant difference in c-di-AMP levels for any of the *nagU* mutants compared to wildtype in the absence of inducer; however, in the presence of inducer c-di-AMP levels decreased significantly for wildtype *nagU* and the inactive variant *nagU*^R106A^, but did not change for the interface-disrupted mutant *nagU*^E26A^ (Fig. 3e. Extended Data Fig. 8b). We also assessed cell size for these strains because previous studies on Δ*gdpP* and *dacA*^G206S^ showed that c-di-AMP levels are negatively correlated with cell size^14, 44^. We found that overexpressing *nagU* or *nagU*^R106A^ significantly increased cell area whereas deleting *nagU* or overexpressing *nagU*^E26A^ did not (Fig. 3f, Extended Data Fig. 8c). Taken together, these results show that NagU can reduce cellular c-di-AMP levels by interacting with DacA. Hence, NagU is a negative regulator of DacA.

### The NagU–DacA interaction sensitizes cells to oxacillin but not vancomycin

The finding that NagU affects c-di-AMP levels when overexpressed led us to ask whether the NagU–DacA interaction influences susceptibility to oxacillin and vancomycin. We showed above that Δ*nagU* cells are hypersensitive to both compounds, and previous studies reported that *S. aureus* cells with low c-di-AMP are hypersensitive to oxacillin whereas cells with high c-di-AMP are protected against oxacillin^44–46^. A spot titer assay confirmed the hypersensitivity of the *dacA* hypomorph (*dacA*^G206S^) to oxacillin (Fig. 3g); we also confirmed that the *ΔgdpP* mutant is protective at higher concentrations of oxacillin (Extended Data Fig. 8d). For vancomycin, we found that *ΔgdpP* was hypersensitive rather than resistant to vancomycin whereas the *dacA* hypomorphic strain was similar to wildtype (Fig. 3g). Hence, *S. aureus* susceptibility to these cell wall antibiotics is differentially affected by c-di-AMP levels.

We next tested effects of the NagU-DacA interaction on antibiotic susceptibility. Under oxacillin treatment, GlmU overexpression was protective, as expected because more UDP-GlcNAc can be made; however, *nagU* overexpression was toxic (Fig. 3h, Extended Data Fig. 8e). To test if the toxicity under *nagU* overexpression was due to the interaction with DacA, we also tested *nagU*^E26A^. This strain was fully viable when overexpressed, implying that the toxicity when wildtype *nagU* was overexpressed resulted from low c-di-AMP levels (Fig. 3h, Extended Data Fig. 8e). For vancomycin, we did not observe toxicity upon overexpressing *nagU*, which is consistent with our finding that low c-di-AMP levels do not influence vancomycin susceptibility (Fig. 3h, Extended Data Fig. 8e). Our findings show that *nagU*-dependent phenotypes for vancomycin are driven only by UDP-GlcNAc synthase activity, whereas for oxacillin both UDP-GlcNAc synthase activity and c-di-AMP levels are important.

### GlmU is not conserved in species having NagU homologues

The discovery that NagU plays an important role in UDP-GlcNAc synthesis in *S. aureus* led us to examine more broadly how NagU is taxonomically distributed across all kingdoms of life. We performed a BLAST search against the Refseq proteins database and identified NagU homologues across eukaryotes and eubacteria (Extended Data Fig. 9a). In eubacteria, NagU is conserved within the family Staphylococcaceae (*e.g., Staphylococcus*, *Mammalicoccus*, *Macrococcus*). It is also commonly found in the PVC superphylum (e.g., *Chlamydiales*) and is sporadically found in *Clostridia* and several other taxa (Fig. 4). The vast majority of NagU orthologues in the family Staphylococcaceae possess a GlmU protein with a defective uridyltransferase domain and we wondered if similar cases exist in other bacterial species carrying NagU homologs. We therefore performed a BLAST search to identify proteins orthologous to full length *S. aureus* GlmU, its N-terminal uridyltransferase or its C-terminal acetyltransferase domain in species having NagU orthologues. We found that the defective GlmU Ser169 variantis only present in Staphylococcaceae (Fig. 4). However, we also found that many bacterial species that possess *nagU* do not appear to encode orthologs of full-length *glmU,* even when the stringency of sequence coverage and identity is reduced (Fig. 4, Extended Data Fig. 9b). Instead, we identified several truncated variations of GlmU. Some species (*e.g., Polystyrenella longa*) curiously have only a truncated N-terminal uridyltransferase domain (Extended Data Fig. 9c), suggesting that there is an unidentified acetyltransferase that can acetylate a glucosamine-phosphate intermediate to complete the pathway. Other species, particularly within the PVC superphylum, encode only a truncated C-terminal acetyltransferase domain (Extended Data Fig. 9d), which likely works with the NagU ortholog to make UDP-GlcNAc. Other organisms (e.g., *Wujia chipingensis*) separately encode truncated forms of each domain, but lack full length GlmU (Extended Data Fig. 9e). Finally, certain bacterial species such as *Clostridium collagenovorans* encode full-length GlmU and both truncated transferase domains (Extended Data Fig. 9f). These findings imply that bacteria beyond Staphylococcaceae combine NagU with multiple variations on GlmU to synthesize UDP-GlcNAc.

**Fig. 4.**
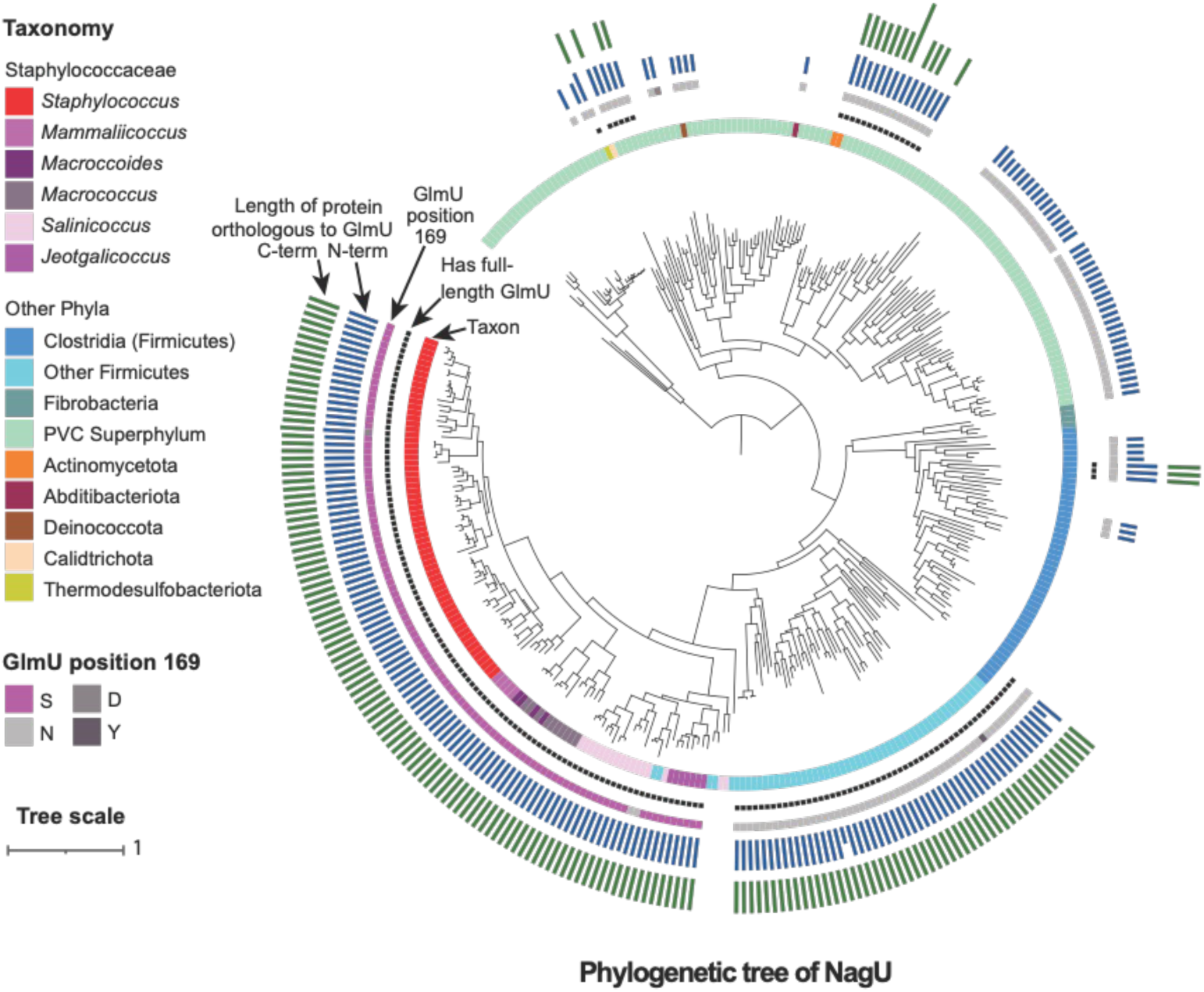
GlmU is not conserved in species having NagU. Phylogenetic tree of NagU proteins from bacterial species representatives. The tree was constructed by aligning hits (>90% coverage and >30% identity) from a *tblastn* search of the Refseq prokaryotic representative genomes database using *S. aureus* NagU as a query. Colored rings, from inside to outside: i) taxonomic distribution of NagU homologs colored by phylum. (Within the Staphylococcaceae, homologs are further colored by genus); ii) the subset of NagU-containing taxa which possess full-length GlmU homologs (>90% coverage and >30% identity; see Extended Data Fig. 9b for lowere stringency GlmU homologs) (black); iii) the identity of the base at GlmU position 169; iv) the relative lengths of *tblastn* hits orthologous to the GlmU N-terminus; and v) the relative lengths of hits orthologous to the GlmU C-terminus, plotted as bars. Phylogenetic visualization was performed using iTOL v7.

## DISCUSSION

The key finding in this study is that *S. aureus* uses a previously uncharacterized monofunctional uridyltransferase called NagU to make most of its UDP-GlcNAc for cell wall synthesis. The uridyltransferase domain of GlmU in *S. aureus* and other Staphylococcaceae is impaired due to an active site mutation. Hence, NagU is essential under cell envelope stress. An evolutionary analysis has shown that NagU orthologues are found in other bacteria, including Chlamydiales and, sporadically, in Clostridia. In a substantial fraction of these organisms, GlmU’s uridyltransferase domain is truncated or missing entirely, and NagU may be the only uridyltransferase that makes UDP-GlcNAc.

An important question arising from this study is why some bacteria rely heavily or exclusively on NagU for UDP-GlcNAc synthesis. One possible answer is that encoding a separate uridyltransferase allows faster production of UDP-GlcNAc. Kinetic analysis of *E. coli* GlmU showed that its acetyltransferase activity is approximately fivefold higher than its uridyltransferase activity. Moreover, GlcNAc-1-P, the product of the acetylation reaction, functions as a feedback inhibitor of the acetylation reaction^13^. Increasing GlmU expression does not get around the rate differential between the two activities of GlmU and so UDP-GlcNAc synthesis will always be rate limiting. However, encoding a uridyltransferase elsewhere in the genome provides a way of circumventing the rate differential bottleneck. More uridyltransferase activity from a separately encoded enzyme may also relieve feedback inhibition at the acetyltransferase step, accelerating overall UDP-GlcNAc biosynthesis. *S. aureus* produces large amounts of peptidoglycan under cell envelope stress as a mechanism to protect cells from lysis and NagU is likely one of the adaptations that allows this.

We have also shown that NagU physically interacts with the c-di-AMP synthase DacA and can negatively regulate its activity. Cyclic-di-AMP, which plays a central role in regulating turgor pressure through control of ion flux across the membrane, is essential for growth in Gram-positive bacteria^15, 40^. Interestingly, NagU is the second peptidoglycan biosynthetic enzyme found to interact with DacA. GlmM, the isomerase that converts GlcN-6-P to GlcN-1-P, also forms a complex with DacA and has been shown to inhibit c-di-AMP synthesis in an *E. coli* expression system^42^. Here, we demonstrated that overexpression of NagU suppresses c-di-AMP synthesis in cells. Although it is unclear why two different peptidoglycan synthesis enzymes would have the ability to negatively regulate DacA, this function is evidently important. We would expect increased expression of either of these enzymes to increase metabolic flux into peptidoglycan precursors, and ultimately into peptidoglycan, and simultaneously to decrease c-di-AMP levels, leading to increased turgor pressure.

Why might there be a link between increased peptidoglycan synthesis and reduced c-di-AMP levels? If the cell wall is thicker, it will be able to withstand higher turgor, but this statement doesn’t explain why higher turgor might be beneficial under conditions where flux into peptidoglycan increases. One possible explanation is that turgor pressure is needed to drive mechanical expansion of the cell wall during growth. The relationship between higher turgor and cell wall expansion during growth has been established in plants, and a recent study in *E. coli* showed that turgor pressure is proportional to the growth rate and directly controls the rate of peptidoglycan synthesis^47, 48^. In this model, turgor pressure drives the outward expansion of the cell wall following cleavage of peptidoglycan crosslinks by autolysins. This expansion creates space to enable newly synthesized peptidoglycan strands and thereby facilitating lateral cell wall growth during cellular expansion.^47^ Such a model could explain the coupling between peptidoglycan synthesis and turgor pressure by NagU: NagU promotes peptidoglycan synthesis by supplying UDP-GlcNAc while simultaneously increasing turgor pressure through inhibition of c-di-AMP synthesis via its physical interaction with DacA (Fig. 5). We note that cells with high c-di-AMP have a thick cell wall but are smaller than wildtype, suggesting that there are differences in how peptidoglycan is incorporated under low and high turgor conditions^44, 49^.

**Fig. 5.**
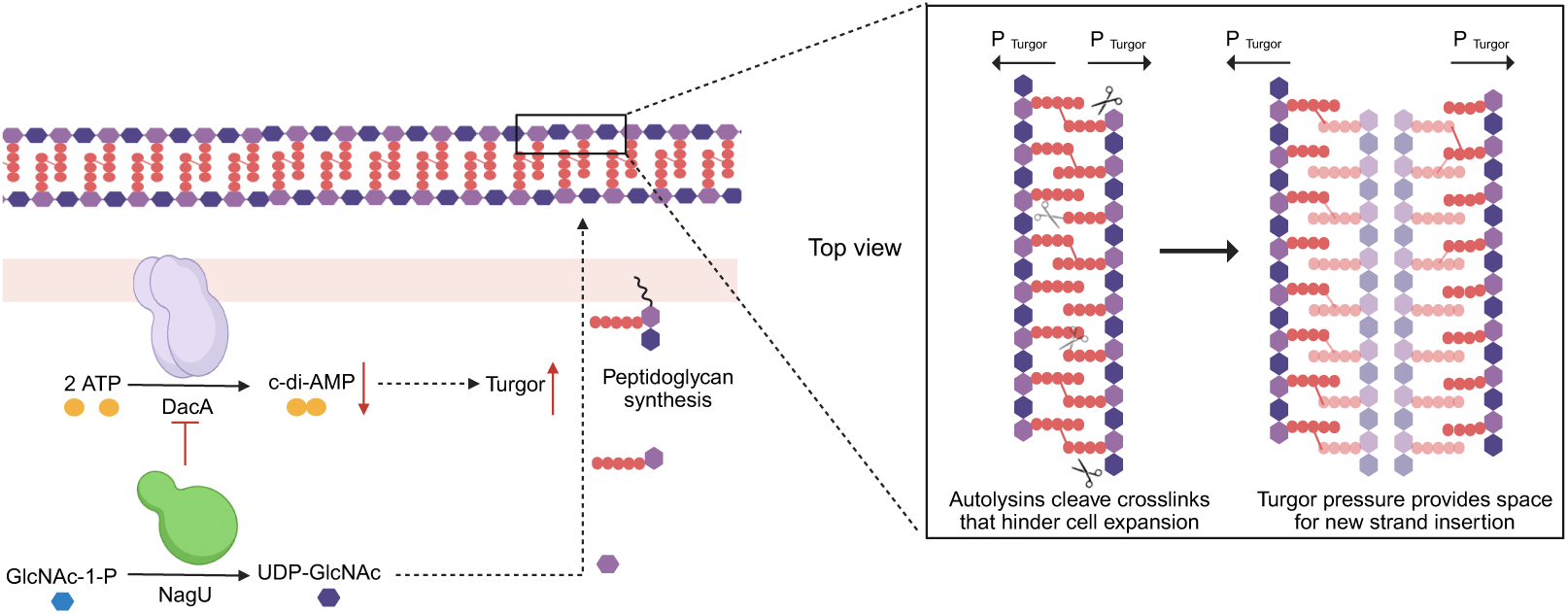
Schematic model illustrating how NagU regulates cell wall synthesis and turgor pressure. Elevated NagU expression enhances flux through the UDP-GlcNAc precursor pool, thereby promoting nascent peptidoglycan synthesis. Concurrently, NagU negatively regulates DacA, the diadenylate cyclase responsible for c-di-AMP production. Reduced intracellular c-di-AMP levels increase cytoplasmic osmolarity, leading to elevated turgor pressure. This rise in turgor pressure drives cell wall expansion by exerting outward force on the existing peptidoglycan, whose crosslinks are cleaved by autolysins. This creates space for the insertion of new peptidoglycan strands for crosslinking, which allows cell wall expansion. The model figure was created using BioRender.

An unanswered question is whether NagU plays any role under DNA damage. The *nagU* operon is upregulated during the SOS response,^23^ but we have found that *nagU* is not protective against ciprofloxacin, which induces the SOS response. Thus, there is currently no evidence that NagU plays a role in the DNA damage response, but it is worth mentioning that there are links between c-di-AMP synthesis and DNA damage induction in other organisms. For example, *Bacillus subtilis* encodes a protein called DisA that makes c-di-AMP and directly senses DNA structure. Upon DNA damage, DisA synthase activity decreases, suggesting it is beneficial to reduce c-di-AMP under this condition^50–52^. Further studies will be required to understand if there is any connection between turgor pressure regulation by NagU and DNA damage repair processes.

We note in closing that the discovery that NagU plays an important role in cell wall biosynthesis in *Staphylococcus aureus* has implications for antibiotic discovery.

## METHODS

### General methods

All *S. aureus* strains used in this study are derived from HG003. Unless otherwise indicated, *S. aureus* strains were cultured at 30 °C in Tryptic Soy Broth (TSB) or on TSB supplemented with 1.5% Bacto agar (TSA). *E. coli* Stellar^TM^ cells (Takara) were used for plasmid construction and BL21(DE3) were used for protein expression. *E. coli* cells were cultured in Lysogeny Broth (LB) or LB supplemented with 1.5% Bacto agar (LBA) except for the purification of DacA_CD_, in which BL21(DE3) cells were grown in TSB. For *S. aureus* strains, antibiotics were used at the following concentrations: 50 μg/mL kanamycin/neomycin (kan/neo), 10 μg/mL erythromycin (erm), and 10 μg/mL chloramphenicol (cam). Anhydrotetracycline (aTc) or isopropyl-β-D-thiogalactopyranoside (IPTG) were used as inducers at 0.4 μM and 1 mM, respectively. For *E. coli* strains, carbenicillin (CB) was used at 100 μg/mL. Strain construction, Tnseq library preparation and Tnseq are described in the Appendix. All strains, plasmids, and oligonucleotides used are listed in Appendix Tables S1–S3.

### RT-qPCR amplification and nucleic acid quantification

Strains were grown in tryptic soy broth (TSB) to mid-log phrase (OD_600_ = ∼0.5) at 30 °C (t0). For SOS-response induction, ciprofloxacin was added to a final concentration of 0.8 µg/mL at t0 and cultures were incubated for an additional 2 h at 30 °C (t1). 1mL of culture was harvested at t0 and t1. Cellular RNA was protected using RNA protection reagent (Qiagen). Cells were lysed using 100 µL TE buffer containing 100 µg/mL lysostaphin for 35 mins at 37 °C. Total RNA was extracted using RNeasy Mini Kit (Qiagen) according to the manufacturer’s protocol. RT-qPCR were performed using the Luna Universal One-Step RT-qPCR kit (New England Biolabs) on a QuantStudio 6 Pro Real-Time PCR System (Applied Biosystems). Gene expression levels were normalized to *rpoB* and relative gene expression was analyzed by the 2^−ΔΔCT^ method.^53^ Results were expressed as relative transcript levels in mutant strains compared with the wild type.

### Spot titers assay

Overnight cultures were adjusted to OD_600_ = 2.0 and diluted 10-fold in TSB to 10^−6^. A 6.5 µL aliquot of each dilution was spotted onto TSA supplemented with the indicated inducers or antibiotics. Plates were incubated at 30 °C for 16 h before imaging. For TSA containing tunicamycin or UgtP inhibitor GLI-2, plates were incubated at 37 °C for 16 h before imaging. For pLOW expression plasmid with erythromycin added, plates were incubated at 30 °C for 40 h before imaging.

### Protein Purification

*S. aureus* HG003 GlmU (SAOUHSC_00471) and mutants (S169N) with N-terminal (6x)-His-tag were expressed from pET-15B in BL21(DE3) *E. coli* grown in LB. Cells were grown in LB medium with 100 µg/mL carbenicillin to OD_600_ = 0.7 – 0.9 at 37 °C and protein expression was induced using 0.25 mM IPTG at 37°C for 3 h. Cells were pelleted and resuspended by 30 mL lysis buffer (50 mM Tris, pH 7.9, 300 mM NaCl, 10 mM imidazole, 5% glycerol, all supplemented with 250 µg/mL DNase I, 5 mM MgCl_2_, 1.0 mg/mL lysozyme, and 1 EDTA free cOmplete™ protease inhibitor cocktail tablet per 50 mL total liquid). Cells were lysed by EmulsiFlex-C3 cell disruptor (Avestin). The protein was affinity-purified using Ni-NTA agarose (Qiagen), washed with increasing concentrations of imidazole and then eluted using 250 mM imidazole. The protein was further purified by Superdex size-exclusion chromatography using a 200 Increase 10/300 GL column (Cytiva) (GE Healthcare) in FPLC buffer (50 mM Tris pH 7.9, 150 mM NaCl, 5% glycerol). The protein was concentrated using a 30 kDa Amicon filter unit (Millipore) and the concentration was determined by BCA assay. Protein was concentrated to 100 μM and stored at −80 °C.

*S. aureus* HG003 DacA_CD_ (SAOUHSC_02407, F101 – K269) with N-terminal (6x)-His-tag is purified in the same way except TSB is used instead of LB. The protein was purified by size-exclusion chromatography using a Superdex 75 Increase 10/300 GL column (Cytiva) (GE Healthcare). The protein was concentrated using a 10 kDa Amicon filter unit (Millipore)

*S. aureus* NagU (SAOUHSC_02423) and mutants (R106A and E26A) with C-terminal (1x)-FLAG tag were purified in the same way with some modifications to the composition of the lysis buffer (1 x TBS with 5% glycerol, all supplemented with 250 µg/mL DNase I, 5 mM MgCl_2_, 1.0 mg/mL lysozyme, and 1 EDTA free cOmplete™ protease inhibitor cocktail tablet per 50 mL total liquid). The protein was purified using Anti-DYKDDDDK G1 Affinity Resin (GenScript) according to manufacturer’s protocol.

### *In vitro* uridylation analysis by HPLC-MS

For heat inactivated NagU, 50 μL 100uM ΝagU-FLAG was heated at 65°C for 15 mins as negative control. *S. aureus* ΝagU-FLAG, its mutant and heated inactivated proteins were added at a final concentration of 1 μM in 10 mM Tris pH 7.4, 100 mM NaCl, and 1 mM MgCl_2_ buffer, substrates GlcNAc-1-P and UTP at a final concentration of 100 μM were added subsequently (Total reaction volume 20 μL) and further incubated at 37°C for 15 min. Reactions were stopped by heating for 5 min at 95 °C and then subsequently cooled to room temperature. 80 μL LCMS-graded acetonitrile were then added. For quantitative analysis, ^13^C_1-_UDP-GlcNAc was spiked in (final concentration 12.5 μM) to calculate the % conversion of the substrate. The reaction was subjected to Agilent 6520 accurate-mass Q-TOF ESI mass spectrometer coupled to an Agilent 1200 series HPLC (Symmetry® C18 5 µm, 3.9 x 150 mm, Waters). The elution method is as follows: flow rate = 0.5 mL/min., 97.0% Solvent A (MS-grade water 97% water, 3% methanol, 10 mM tributylamine, 9.8 mM acetic acid) 3% Solvent B (MS-grade methanol) for 3 minutes, followed by a linear gradient of Solvent B from 3.0% to 100% over 30 minutes, then 5 minutes of 100% Solvent B. The following settings were used for the mass spectrometer: negative ion mode; mass range of 200 to 1000 m/z; drying gas: 325 °C, 7 L/min.; nebulizer: 50 psig; capillary: 3500 V; fragmentor: 150 V; skimmer: 65 V; octupole 1 RF Vpp: 750 V; 125 ms/spectrum). For quantitative analysis, the peak areas for the mass signal corresponding to UDP-GlcNAc were integrated for each sample separately, and then normalized to the internal standard ^13^C_1_-UDP-GlcNAc. The average values of biological triplicates and SEM were plotted.

### Transmission electron microscopy

Overnight cultures of the indicated strains were diluted to OD_600_ = 0.05 followed by the addition of 1 mM IPTG. The samples were further cultured for 3 h at 30 °C before the addition of equal volume of fixatives (2.5% paraformaldehyde, 5.0% glutaraldehyde, 0.06% picric acid in 0.2 M cacodylate buffer). The samples were sent and processed The Harvard Medical School Electron Microscopy Facility and were imaged by JEOL 1200EX. Cell wall thickness was quantified manually directly form the results image by measuring the widths of the cell wall section in the image. Cell wall thickness was measured at the cell periphery.

### Targeted metabolomics

The targeted metabolomics assay is modified from a previously reported method^14^. Briefly, overnight cultures of *S. aureus* cells were diluted to 0.05 in TSB (20 mL) and incubated for 5 h at 30 °C. For overexpression, IPTG was added to a final concentration of 1 mM. A 10 mL aliquot was centrifuged at 9,000×*g* for 5 min, washed by 1x TBS and lyophilized to determine the dry weight for normalization purposes. A 5 ml aliquot from the same culture was centrifuged at 9,000×*g* for 5 min. The pellet was suspended in 1 mL ice-cold extraction buffer (LC-MS grade acetonitrile/methanol/water 2∶2∶1 containing 0.25 µM ^15^N_10_-c-di-AMP (BIOLOG Life Science Institute) as internal standard). Samples were snap frozen with liquid nitrogen for 15 s before being heated to 95 °C for 10 min. Samples were mixed with 0.5 ml of 0.1 mm glass beads and lysed in a Fast-Prep-24 Classic bead beating grinder and lysis system for 45 sec at setting 2 twice (MP Biomedicals). Glass beads were separated by centrifugation at 17,000×*g* for 5 min at 4°C. The supernatant was removed and stored at 4°C and the remaining glass beads/cell debris mixture was washed with 1 ml extraction buffer without internal standard, incubated on ice for 15 min and again lysed. Samples were centrifuged again and the supernatant combined with the previous one. Glass beads were washed again with 1 ml extraction buffer, incubated on ice for 15 min and centrifuged. All supernatant were combined and 3 mL LCMS graded water was added before lyophilization. 200 μL water was added and samples were subject to the UPLC-MS.

UPLC-MS analysis was performed on a Vanquish UHPLC coupled with Orbitrap Exploris 240 high-resolution MS system (Thermo Fisher Scientific) equipped with Waters ACQUITY^TM^ Premier HSS T3 100 Å, 1.8 µm, 2.1 × 100 mm column for the analysis of GlcNAc-1-P. Waters Atlantis^TM^ Premier BRH Z-HILIC T3 100 Å, 1.7 µm, 2.1 × 100 mm column was used for the analysis of c-di-AMP. For the analysis of GlcNAc-1-P, the elution method is as follows: flow rate = 0.3 mL/min starting from 1 % Solvent A (LCMS-grade 10mM ammonium bicarbonate) and 99% Solvent B (LCMS-grade 90% acetonitrile 10mM ammonium bicarbonate), followed by a linear gradient of Solvent B from 99% to 40% over 10 minutes, then 1 minute of 40% solvent B followed by a linear gradient of Solvent B from 40% to 99% over 1 minute. The column was then equilibrated with 99% Solvent B for 9 mins. The following settings were used for the mass spectrometer: positive ion mode; mass range: 120 to 1300 m/z; Spray voltage: 3500V; Sheath Gas: 50 Arb; Aux Gas: 10 Arb; Sweep Gas: 1 Arb; Ion transfer tuber temperature: 325 °C; Vaporizer temperature: 350 °C. For the analysis of c-di-AMP, the elution method is as follow: flow rate = 0.3 mL/min., Solvent A (LCMS-grade 10mM ammonium formate, 0.1% formic acid) for 2.5 mins, followed by a linear gradient of Solvent B (LCMS-grade methanol) from 3.0% to 30% over 8 minutes, then 2 minutes of 100% Solvent A. The following settings were used for the mass spectrometer: negative ion mode; mass range: 120 to 1300 m/z; Spray voltage: 3000V; Sheath Gas: 50 Arb; Aux Gas: 10 Arb; Sweep Gas: 1 Arb; Ion transfer tuber temperature: 325 °C; Vaporizer temperature: 350 °C. The peak areas for the mass signal corresponding to these metabolites were integrated for each sample separately, and then normalized to internal standard ^15^N_10_-c-di-AMP and then further normalized to dry mass and WT. The average values of biological triplicate and SEM were plotted.

### Enzymatic assays

Kinetic parameters for NagU-FLAG and His_6_-GlmU is determined using the EnzChek™ Pyrophosphate Assay Kit (Invitrogen) according to the manufacturer protocol^54^. Product formation was monitored at λ = 360 nm using a Spectramax microplate reader (Molecular Devices). To determine *K*_m_, the concentration of GlcNAc-1-P or UTP was varied from 7.8 to 500 µM at a fixed concentration of 500 µM UTP or 500 µM GlcNAc-1-P, respectively. The enzyme concentration was 5 ng/mL.

### Sequence Logo construction for GlmU

An initial sequence list of GlmU orthologs was acquired from EV couplings server using GlmU (SAOUHSC_00471) sequence as queries (Sequence database: UniRef90)^55^. Sequences were downloaded and the taxonomic lineage information was acquired using UniProt ID mapping to obtain UniProtKB entries. Sequences from unclassified bacteria were trimmed and in total 6395 proteins were identified and 40 of these were from Staphylococcaceae (Supplementary Table 4). Sequence were aligned with Clustal Omega 1.2.3. Selective regions of aligned sequence were made using WebLogo^56^.

### Immunoprecipitation assays of FLAG-tagged proteins from *S. aureus*

The immunoprecipitation assay was performed accordingly to previously established protocol^57^. Strains LF1148, LF1143, LF1150 and LF1159 were used.

### AlphaFold2 Modeling of DacA-NagU complexes

Predicted AlphaFold2-based models of DacA-NagU complexes was generated using ColabFold v1.5.5^58^. Modeling was performed by template null AlphaFold2 multimer v3 with 20 recycles. The multiple sequence alignment (MSA) mode was “mmseqs2_uniref_env” with unpaired and paired mode. Complexes were visualized in UCSF ChimeraX 1.10^59^. Output files for all AlphaFold2 models are available with this manuscript online (Supplementary Data 1).

### Size exclusion chromatography with multi-angle light scatterings

To assemble the complex, purified NagU-FLAG and His_6_-DacA_CD_ were mixed at the indicated mass ratio and incubated overnight at 4 °C. The mixture was fractionated by SEC on a Superdex 75 Increase 10/300 GL column (Cytiva) equilibrated in HBS (25 mM HEPES pH 7.5, 150 mM NaCl). Peak fractions corresponding to the putative complex were pooled and concentrated to 1 mg mL^−1^ using a 30 kDa MWCO centrifugal concentrator (Amicon). Samples were centrifuged at 20,000 rcf for 1 min to remove any aggregates and injected at 100 µL (1 mg mL^−1^) onto a Superdex 200 Increase 10/300 GL column (Cytiva) for samples with excess NagU-FLAG and SRT^®^ SEC-150 (5 µm) HPLC columns (Supelco) for sample with excess His_6_-DacA_CD_ equilibrated in HBS at room temperature. The flow rate was 0.5 mL min^−1^. The SEC was coupled in-line to a multi-angle light scattering detector (DAWN HELEOS II, Wyatt Technology), an Optilab differential refractive index (dRI) detector (Wyatt), and a UV detector (λ = 280 nm). Instrument alignment/normalization and inter-detector delay volumes were calibrated immediately before sample runs using bovine serum albumin (BSA; 2 mg mL^−1^, 100 µL injection). Light-scattering and dRI signals were analyzed with ASTRA software (Wyatt Technology). The protein refractive index increment was set to dn/dc = 0.185 mL g⁻¹, and extinction coefficients were calculated from amino-acid composition. Absolute, weight-average molar weight (MW) across elution peaks were obtained using the Zimm formalism, assuming isotropic scatterers. Monodispersity was assessed from the Mw distribution across each peak.

### Wide-field epifluorescence microscopy and cell area quantification

Strains were normalized to OD_600_ = 0.05 in TSB with 1 mM IPTG, 5 μg/mL trimethoprim, and 5 μg/mL erythromycin, and grown at 30 °C to early log phase (OD_600_ = ∼0.3). Nile red was added to a final concentration of 5 μg/mL and incubated at 37 °C for 5 minutes. Cells were spotted onto TSB pads containing 2% agarose for imaging. Cells were imaged using brightfield, phase-contrast, and wide-field epifluorescence microscopy at the MicRoN (Microscopy Resources on the North Quad) facility, Harvard Medical School. Images were obtained using a Nikon Ti inverted microscope fitted with a custom-made cage incubator set at 30°C, a Nikon motorized stage with an OkoLab gas incubator and a slide insert attachment, an Andor Zyla 4.2 Plus sCMOS camera, Lumencore SpectraX LED Illumination, Plan Apo lambda 100x/1.45 Oil Ph3 DM objective lens, and Nikon Elements 4.30 acquisition software. The microscope was fitted with Chroma ET filter cubes in a motorized filter turret: GFP (49002) and mCherry (49008). The following exposure times were used: 50ms (Nile red labelling), 500 ms (cytoplasmic mNeonGreen).

Phrase contrast and fluorescent images were analyzed in FIJI (version 2.16.0/1.54p) using the MicrobeJ plugin (version 15.3p)^60, 61^. To remove artifactual halos around cells in phase contrast images and improve segmentation fidelity, phase contrast image pixel values were inverted, then subjected to rolling ball (25-pixel diameter) dark background subtraction, then pixel values inverted again. Fluorescent images were subjected to rolling ball (50-pixel diameter) dark background subtraction. In MicrobeJ, cell segmentation was performed on background-subtracted phase contrast images with the default parameters unless otherwise noted: Bright background, Otsu thresholding, offset = 11, area minimum = 0.5 μm^2^, exclude on edges selected, segmentation selected. Cells were removed or split manually as required, informed by the Nile Red and cytoplasmic mNeonGreen signals to resolve small clumps. At least 500 cells per biological replicate were used for cell area measurements. Resulting measurements were visualized as Violin SuperPlots^62^, and the mean of means compared between experimental strains and the control strain using a one-way ANOVA followed by Tukey’s tests with Bonferroni corrections.

### Comparative genomics

Using *S. aureus* amino acid sequences for NagU (SAOUHSC_02423) and GlmU (SAOUHSC_00471), three BLAST databases (all downloaded July 28, 2025) were searched using thresholds of 30% identity and 90% coverage of the query protein: i) the Refseq Protein database using blastp; ii) the Refseq prokaryotic genomes database using tblastn; and iii) all 2,424 high-quality Refseq genomes for *Staphylococcus*, using tblastn. The NCBI taxonomy database (downloaded July 28, 2025) was used to classify the taxonomic distribution of hits^63^. To generate protein trees of Blast hits, amino acid sequences were aligned using the Muscle aligner^64^, and trees were generated using FastTree^65^. Itol was used to visualize trees^66^. Muscle alignments were used to determine the presence of specific mutations (GlmU Ser169) in each sequence.

To assess the taxonomic distribution of individual domains within the GlmU protein, BLAST searches were performed separately using the *S. aureus* GlmU uridyltransferase and acetyltransferase (amino acids 1-251 and 251-442, respectively), using coverage thresholds ranging from 70% to 90% and identity thresholds between 25% and 30%.

## Supporting information

Supplementary Table 2 Foldseek result

Supplementary Table 5 Protein MS

Supplementary Table 1 Tn-seq result

Supplementary Table 3 Kinetic parameters of NagU and GlmU

Supplementary Table 4 GlmU sequence alignment

## ACKNOWLEDGEMENTS

We thank the staff at the Analytical Chemistry Core at Harvard Medical School for providing access to liquid chromatography-mass spectrometry instrumentation. We thank the staff at the Electron Microscopy Facility at Harvard Medical School for Electron Microscopy Imaging, consultation and services. We gratefully acknowledge the MicRoN (Microscopy Resources on the North Quad) Core at Harvard Medical School for their support and assistance in the fluorescent microscopy. We thank the Center for Macromolecular Interactions at Harvard Medical School for support and assistance in SECMALS analysis. We thank thew Taplin Biological Mass Spectrometry Facility for mass spectrometry analysis of proteins. Funding for this work was provided by the NIH through P01 (AI083214) and R01 (AI148752).

## AUTHOR CONTRIBUTION

P-M. F. and S.W designed research; P-M. F., A. W., A. M., Y. P performed the experiments and analysis; S.W and A.E supervised the study; P-M. F. and S.W wrote the paper.

---

**Extended Data Fig. 1.**
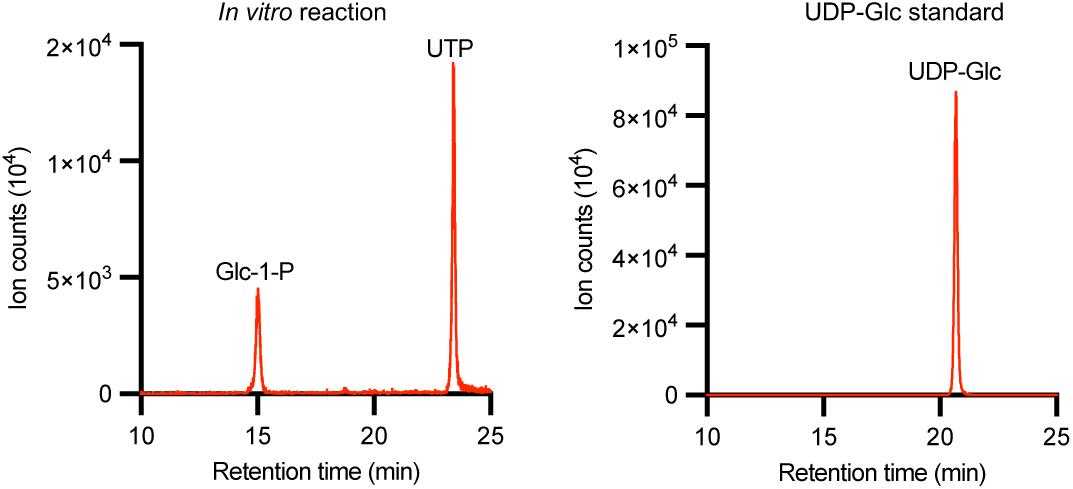
NagU cannot synthesize UDP-Glucose from glucose-1-phosphate. LC-MS chromatograms of the attempted *in vitro* NagU-mediated uridyltransfer reaction using glucose-1-phosphate (Glc-1-P) and UTP as substrates (Left). Glucose-1-phosphate (Glc-1-P) and UTP were used as substrate (Left). The LC trace of a UDP-Glucose (UDP-Glc) standard is shown for comparison (Right).

**Extended Data Fig. 2.**
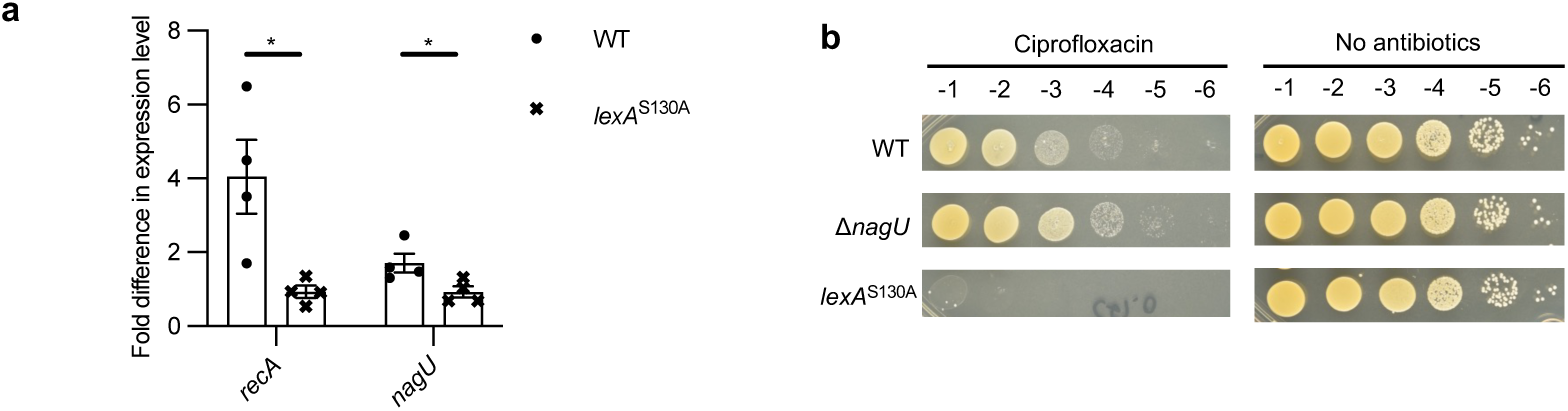
The expression of *nagU* increases upon the addition of ciprofloxacin but Δ*nagU* does not potentiate ciprofloxacin. **a**, The fold change detected by RT-qPCR in expression level of *nagU* of HG003 WT and *lexA*^S130A^ after the addition of 0.8 μg mL⁻¹ ciprofloxacin for 2 hr. *recA* acts as the positive control. Data are means ± SEM of 4 biological replicates. \**P* < 0.05. Statistical significance was determined by two-tailed unpaired Student *t*-test. **b**. Spot titer assays showing that the *lexA*^S130A^ rather than Δ*nagU* hypersensitizes cells to ciprofloxacin. Strains spotted include WT HG003, Δ*nagU* and *lexA*^S130A^. Strains were spotted on TSA with (Left) or without (Right) 0.125 μg mL⁻¹ ciprofloxacin.

**Extended Data Fig. 3.**
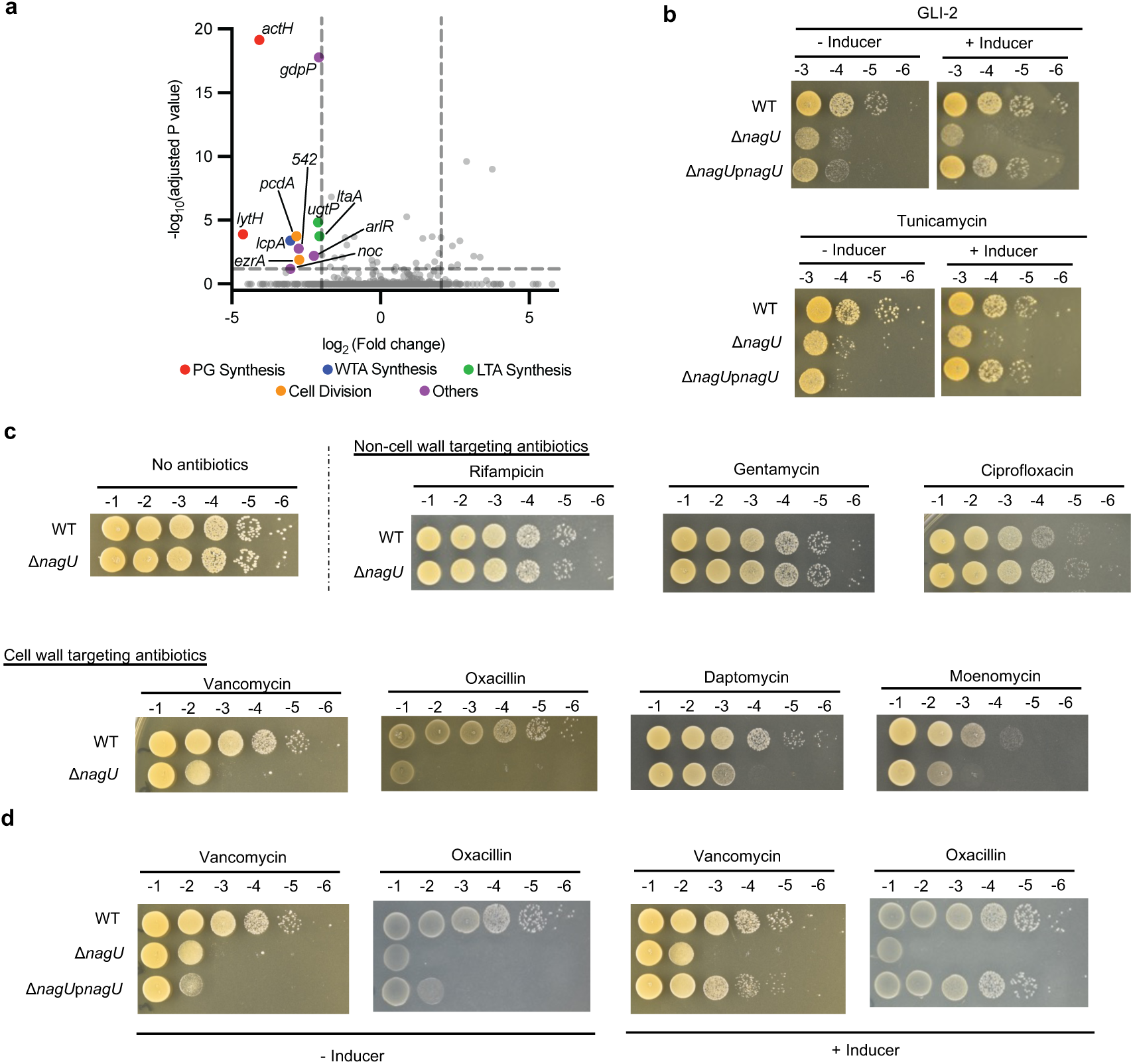
Deletion of *nagU* sensitizes the cells to cell wall targeting antibiotics. **a**, Volcano plot of transposon insertions within individual genes in a Δ*nagU* transposon library relative to a WT control library. Significantly depleted genes (>4-fold, adjusted *P* < 0.05) are labelled. **b**, Spot titer assays of WT, Δ*nagU*, and the *nagU*-complemented strain on TSA containing 2.0 μg ml⁻¹ GLI-2 (Top) or 0.2 μg mL⁻¹ tunicamycin (Bottom). Inducer 0.4 µM anhydrotetracycline was added for complementation. **c**, Spot titer assays of WT and Δ*nagU* strains across a panel of antibiotics, including 1 ng mL⁻¹ rifampicin, 1.25 µg mL⁻¹ gentamicin, 0.125 µg mL⁻¹ ciprofloxacin, 0.725 µg mL⁻¹ vancomycin, 0.1 µg mL⁻¹ oxacillin, 1.875 µg mL⁻¹ daptomycin, and 12.5 ng mL⁻¹ moenomycin. **d**, Spot titer assays of WT, Δ*nagU*, and the *nagU*-complemented strain on TSA containing 0.725 µg mL⁻¹ vancomycin or 0.1 µg mL⁻¹. Inducer = 0.4 µM anhydrotetracycline.

**Extended Data Fig. 4.**
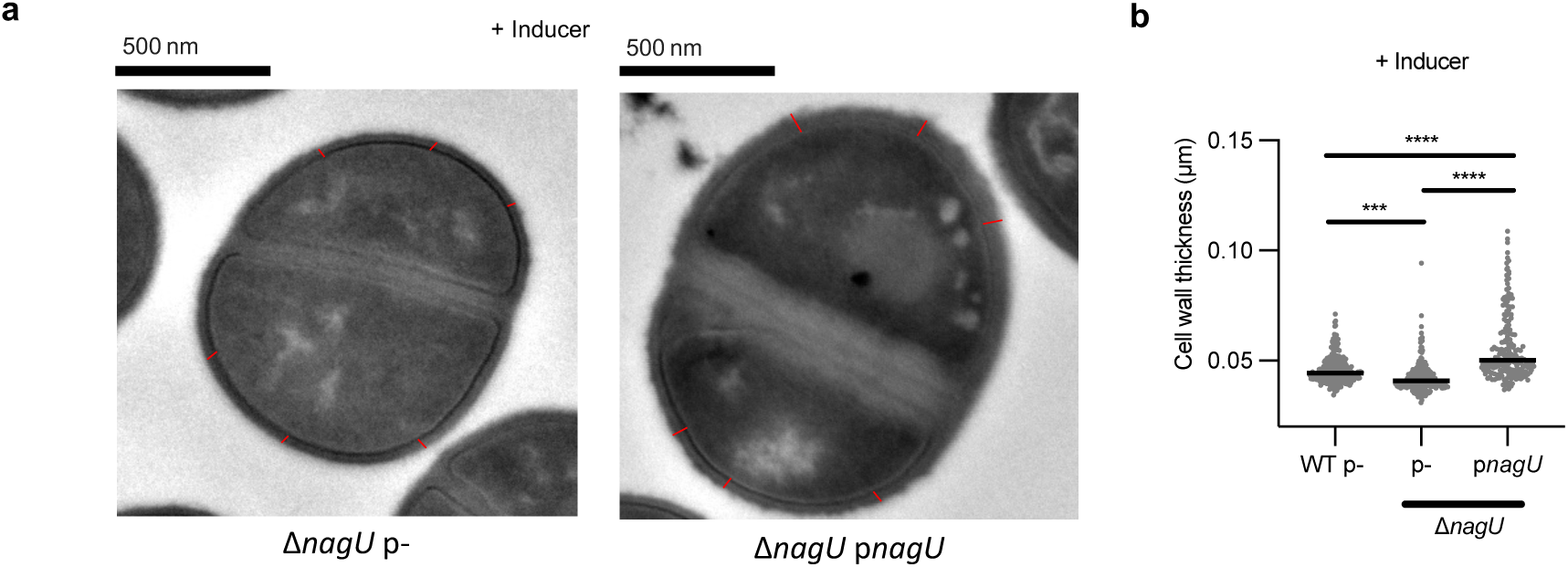
*nagU* overexpression increases cell wall thickness. **a**, Representative transmission electron micrographs of a Δ*nagU* cell carrying empty vector or a *nagU* expressing vector. Inducer 1 mM IPTG was added for overexpression. Red lines indicate cell walls thickness are measured at poles or peripheries (Average of 6 – 10 measurements per cell for statistical analysis in Extended Data Fig. 4b). Scale bars, 500 nm. **b**, Quantification of cell wall thickness across wildtype carrying empty vector, Δ*nagU* carrying empty vector or a *nagU* expressing vector. Inducer 1 mM IPTG was added for overexpression. Total cell counts: 235 (WT p-), 220 (Δ*nag* p-), and 191 (Δ*nag* p*nagU*). \*\*\**P* < 0.001, \*\*\*\**P* < 0.0001. Statistical significance was determined by one-way ANOVA with Dunnett correction. Data are representative of two independent experiments.

**Extended Data Fig. 5.**
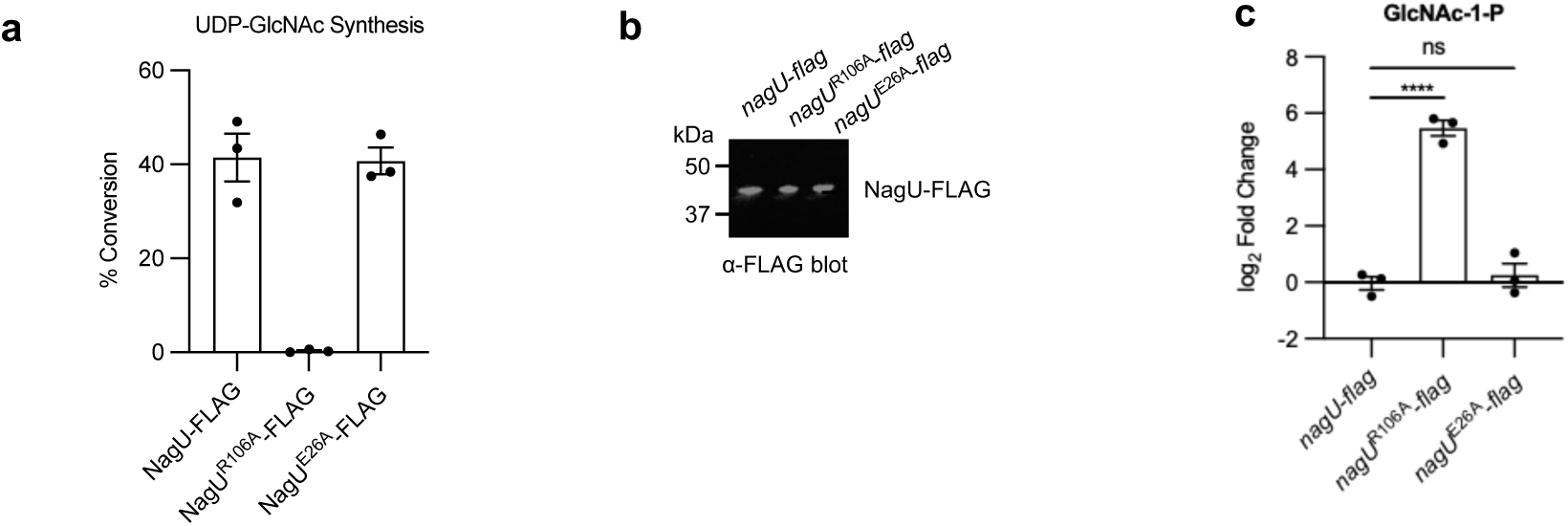
NagU^R106A^ is a catalytic inactive variant and NagU^E26A^ is a NagU-DacA interface-disrupted variant. **a***. In vitro* uridyltransfer assays for UDP-GlcNAc synthesis using NagU-FLAG, NagU^R106A^-FLAG, and NagU^E26A^-FLAG. NagU^R106A^-FLAG. **b**. Western blot analysis of NagU-FLAG variants expressed from the native chromosomal locus in wildtype *nagU*^R106A^ and *nagU*^E26A^ strains. **c**. UPLC measurement of cellular GlcNAc-1-P levels in wildtype, *nagU*^R106A^ and *nagU*^E26A^. Data are means ± SEM of 3 biological replicates. \*\*\*\**P* < 0.0001. ns, not significant. Statistical significance was determined by one-way ANOVA with Dunnett correction.

**Extended Data Fig.6.**
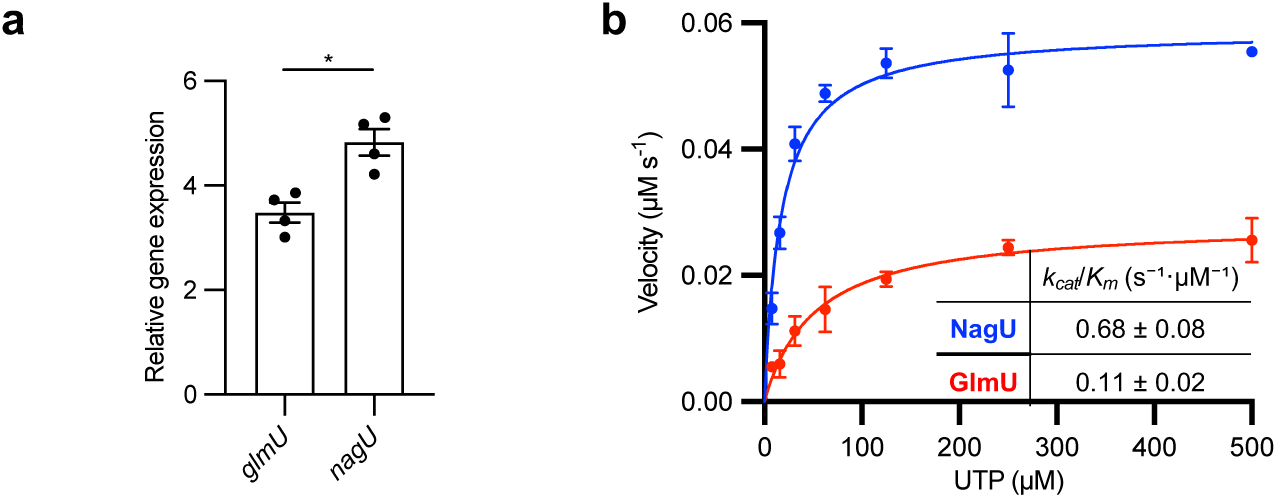
NagU is a primary uridyltransferase. **a.** Relative expression of *glmU* and *nagU* in early log-phase wildtype HG003, detected by RT-qPCR and normalized to *rpoB*. Data are means ± SEM of 4 biological replicates. \**P* < 0.05. Statistical significance was determined by two-tailed paired Student *t* test. **b**. Michaelis-Menten kinetics of NagU and GlmU using UTP as substrate. *k_cat_*/*K_m_* are means ± SEM of 3 biological replicates (See Supplementary table 3).

**Extended Data Fig 7.**
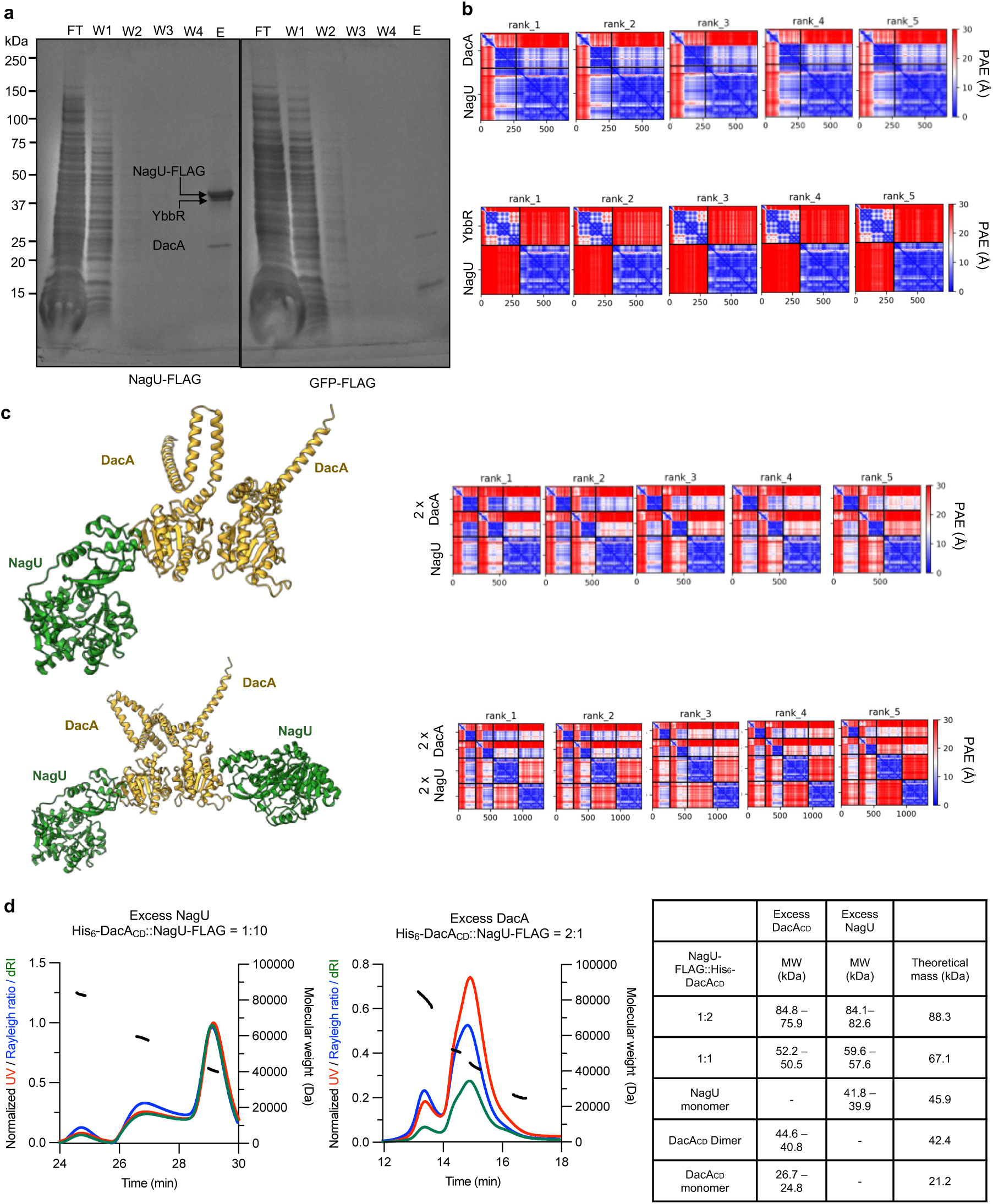
NagU physically interacts with DacA. **a**, Pull-down assays identifying DacA-YbbR complex as the interaction partners of NagU. NagU-FLAG (Left) or GFP-FLAG (Right) were used as the bait. **b**. Position alignment error (PAE) plots from ColabFold for NagU-DacA (Top) and NagU-YbbR (Bottom) structural predictions. **c.** PAE plots for ColabFold-predicted NagU:DacA complexes at 1:2 and 2:2 stoichiometries. **d.** SEC-MALs chromatographs of NagU:DacA complex formation under conditions of excess NagU (Left) or excess DacA_CD_ (Middle). Experimental and theoretical mass of the complexes were depicted in table (Right).

**Extended Data Fig. 8.**
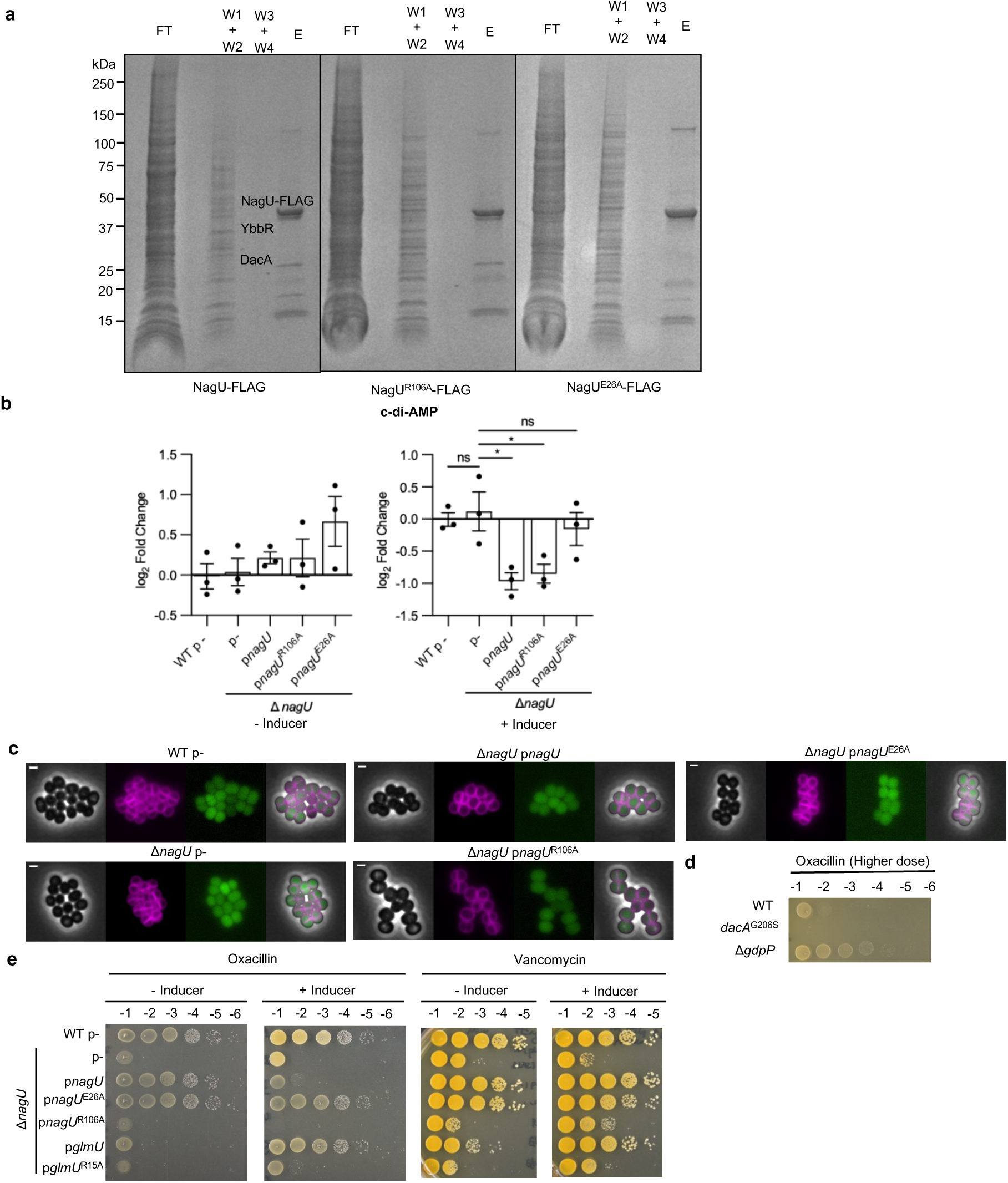
NagU^E26A^ mutation disrupts the physical interaction between DacA and NagU. **a**, Pull-down assays testing the interaction between the DacA-YbbR complex and NagU-FLAG (Left) and NagU^R106A^ -FLAG (Middle) and NagU^E26A^-FLAG (Right)as bait. **b,** UPLC quantification of c-di-AMP levels upon the overexpression of *nagU* variants of *glmU* variants. Strains spotted include wildtype carrying empty vector, Δ*nagU* carrying empty vector or vector expressing *nagU*, *nagU*^R106A^*, nagU*^E26A^*, glmU* and *glmU*^R15A^. Strains were cultured in TSA with 10 µg mL⁻¹ erythromycin. Inducer 1 mM IPTG was added for overexpression. Data are means ± SEM of 3 biological replicates. \*\*\**P* < 0.001, \**P* < 0.05, ns, not significant; Statistical significance was determined by one-way ANOVA with Dunnett corrections. **c,** Representative fluorescent images of strains WT carrying empty vector, Δ*nagU* carrying empty vector or vector expressing *nagU*, *nagU*^R106A^ and*, nagU*^E26A^. (From left: brightfield, nile red, GFP, overlay). Scale bar: 1 µm. **d**. Spot titer of WT, *dacA*^G206S^ and Δ*gdpP* on TSA plates containing 0.3 µg mL⁻¹ oxacillin. **e.** Spot titer comparing the effect of *nagU* and *glmU* variant on vancomycin and oxacillin sensitivity. Strains spotted include WT carrying empty vector, Δ*nagU* carrying empty vector or vector expressing *nagU*, *nagU*^R106A^*, nagU*^E26A^*, glmU* and *glmU*^R15A^. Strains are spotted on TSA containing 0.325 µg mL⁻¹ vancomycin or 0.1 µg mL⁻¹ oxacillin. Inducer 1 mM IPTG was added for overexpression.

**Extended Data Fig. 9.**
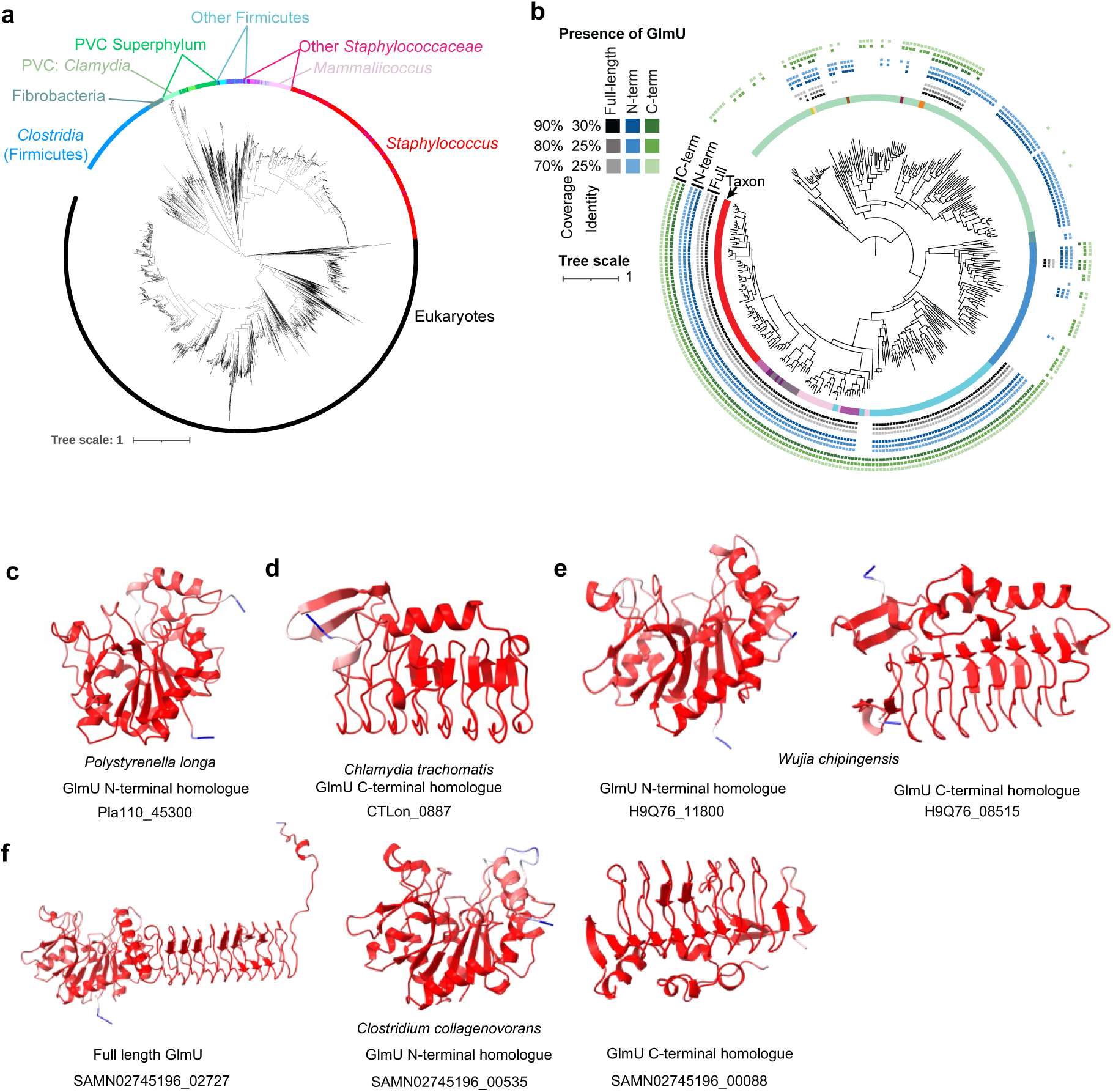
GlmU is not conserved for the bacteria having NagU homologues. **a,** Phylogenetic tree of NagU proteins from representative bacterial species. This tree was constructed as in Figure 4. Colored rings, from inside to outside, indicate i, taxonomic distribution of NagU homologs colored by phylum, as in Figure 4. ii–iv, presence of full-length GlmU; v–vii, presence of the GlmU N-terminal domain; viii–x, presence of the GlmU C-terminal domain across three BLAST thresholds. Phylogenetic visualization was performed in iTOL v7. **c-f,** Alphafold predicted structure of homologues of full-length GlmU, truncated GlmU with N-terminus only and GlmU with C-terminus only in different bacterial species colored by pLDDT confidence score (Blue: low, Red: high). Structures were obtained from AlphaFold Protein Structure Database: Pla110_45300 (A0A518CU88), CTLon_0887 (A0A6H2W3I8), H9Q76_11800 (A0A7G9FLF2), H9Q76_08515 (A0A7G9FJQ8), SAMN02745196_02727 (A0A1M5Y8K2), SAMN02745196_00535 (A0A1M5TDI8) and SAMN02745196_00088 (A0A1M5S9K9).

## REFERENCES

(1) Silhavy, T. J.; Kahne, D.; Walker, S. The bacterial cell envelope. Cold Spring Harb Perspect Biol 2010, 2 (5), a000414. DOI: 10.1101/cshperspect.a000414.

(2) Vollmer, W.; Blanot, D.; de Pedro, M. A. Peptidoglycan structure and architecture. FEMS Microbiol Rev 2008, 32 (2), 149–167. DOI: 10.1111/j.1574-6976.2007.00094.x.

(3) Puls, J. S.; Brajtenbach, D.; Schneider, T.; Kubitscheck, U.; Grein, F. Inhibition of peptidoglycan synthesis is sufficient for total arrest of staphylococcal cell division. Sci Adv 2023, 9 (12), eade9023. DOI: 10.1126/sciadv.ade9023.

(4) Wang, M.; Buist, G.; van Dijl, J. M. Staphylococcus aureus cell wall maintenance – the multifaceted roles of peptidoglycan hydrolases in bacterial growth, fitness, and virulence. FEMS Microbiol Rev 2022, 46 (5). DOI: 10.1093/femsre/fuac025.

(5) Culp, E. J.; Waglechner, N.; Wang, W.; Fiebig-Comyn, A. A.; Hsu, Y. P.; Koteva, K.; Sychantha, D.; Coombes, B. K.; Van Nieuwenhze, M. S.; Brun, Y. V.;, et al. Evolution-guided discovery of antibiotics that inhibit peptidoglycan remodelling. Nature 2020, 578 (7796), 582–587. DOI: 10.1038/s41586-020-1990-9.

(6) Lewis, K.; Lee, R. E.; Brotz-Oesterhelt, H.; Hiller, S.; Rodnina, M. V.; Schneider, T.; Weingarth, M.; Wohlgemuth, I. Sophisticated natural products as antibiotics. Nature 2024, 632 (8023), 39–49. DOI: 10.1038/s41586-024-07530-w.

(7) Page, J. E.; Walker, S. Natural products that target the cell envelope. Curr Opin Microbiol 2021, 61, 16–24. DOI: 10.1016/j.mib.2021.02.001.

(8) Sarkar, P.; Yarlagadda, V.; Ghosh, C.; Haldar, J. A review on cell wall synthesis inhibitors with an emphasis on glycopeptide antibiotics. Medchemcomm 2017, 8 (3), 516–533. DOI: 10.1039/c6md00585c.

(9) Bugg, T. D.; Braddick, D.; Dowson, C. G.; Roper, D. I. Bacterial cell wall assembly: still an attractive antibacterial target. Trends Biotechnol 2011, 29 (4), 167–173. DOI: 10.1016/j.tibtech.2010.12.006.

(10) Olsen, L. R.; Roderick, S. L. Structure of the *Escherichia coli* GlmU pyrophosphorylase and acetyltransferase active sites. Biochemistry 2001, 40 (7), 1913–1921. DOI: 10.1021/bi002503n.

(11) Mochalkin, I.; Lightle, S.; Zhu, Y.; Ohren, J. F.; Spessard, C.; Chirgadze, N. Y.; Banotai, C.; Melnick, M.; McDowell, L. Characterization of substrate binding and catalysis in the potential antibacterial target N-acetylglucosamine-1-phosphate uridyltransferase (GlmU). Protein Sci 2007, 16 (12), 2657–2666. DOI: 10.1110/ps.073135107.

(12) Mengin-Lecreulx, D.; van Heijenoort, J. Identification of the glmU gene encoding N-acetylglucosamine-1-phosphate uridyltransferase in Escherichia coli. J Bacteriol 1993, 175 (19), 6150–6157. DOI: 10.1128/jb.175.19.6150-6157.1993.

(13) Mengin-Lecreulx, D.; van Heijenoort, J. Copurification of glucosamine-1-phosphate acetyltransferase and *N*-acetylglucosamine-1-phosphate uridyltransferase activities of *Escherichia coli*: characterization of the glmU gene product as a bifunctional enzyme catalyzing two subsequent steps in the pathway for UDP-*N*-acetylglucosamine synthesis. J Bacteriol 1994, 176 (18), 5788–5795. DOI: 10.1128/jb.176.18.5788-5795.1994.

(14) Corrigan, R. M.; Abbott, J. C.; Burhenne, H.; Kaever, V.; Grundling, A. c-di-AMP is a new second messenger in *Staphylococcus aureus* with a role in controlling cell size and envelope stress. PLoS Pathog 2011, 7 (9), e1002217. DOI: 10.1371/journal.ppat.1002217.

(15) Brogan, A. P.; Bardetti, P.; Rojas, E. R.; Rudner, D. Z. Cyclic-di-AMP modulates cellular turgor in response to defects in bacterial cell wall synthesis. Nat Microbiol 2025, 10 (7), 1698–1710. DOI: 10.1038/s41564-025-02027-2.

(16) Foster, A. J.; van den Noort, M.; Poolman, B. Bacterial cell volume regulation and the importance of cyclic di-AMP. Microbiol Mol Biol Rev 2024, 88 (2), e0018123. DOI: 10.1128/mmbr.00181-23.

(17) Barreteau, H.; Kovac, A.; Boniface, A.; Sova, M.; Gobec, S.; Blanot, D. Cytoplasmic steps of peptidoglycan biosynthesis. FEMS Microbiol Rev 2008, 32 (2), 168–207. DOI: 10.1111/j.1574-6976.2008.00104.x.

(18) Mio, T.; Yamada-Okabe, T.; Arisawa, M.; Yamada-Okabe, H. Saccharomyces cerevisiae GNA1, an essential gene encoding a novel acetyltransferase involved in UDP-N-acetylglucosamine synthesis. J Biol Chem 1999, 274 (1), 424–429. DOI: 10.1074/jbc.274.1.424.

(19) Milewski, S.; Gabriel, I.; Olchowy, J. Enzymes of UDP-GlcNAc biosynthesis in yeast. Yeast 2006, 23 (1), 1–14. DOI: 10.1002/yea.1337 From NLM Medline.

(20) Mio, T.; Yabe, T.; Arisawa, M.; Yamada-Okabe, H. The eukaryotic UDP-*N*-acetylglucosamine pyrophosphorylases. Gene cloning, protein expression, and catalytic mechanism. J Biol Chem 1998, 273 (23), 14392–14397. DOI: 10.1074/jbc.273.23.14392.

(21) van Kempen, M.; Kim, S. S.; Tumescheit, C.; Mirdita, M.; Lee, J.; Gilchrist, C. L. M.; Soding, J.; Steinegger, M. Fast and accurate protein structure search with Foldseek. Nat Biotechnol 2024, 42 (2), 243–246. DOI: 10.1038/s41587-023-01773-0.

(22) Grundling, A.; Schneewind, O. Genes required for glycolipid synthesis and lipoteichoic acid anchoring in Staphylococcus aureus. J Bacteriol 2007, 189 (6), 2521–2530. DOI: 10.1128/JB.01683-06.

(23) Cirz, R. T.; Jones, M. B.; Gingles, N. A.; Minogue, T. D.; Jarrahi, B.; Peterson, S. N.; Romesberg, F. E. Complete and SOS-mediated response of *Staphylococcus aureus* to the antibiotic ciprofloxacin. J Bacteriol 2007, 189 (2), 531–539. DOI: 10.1128/JB.01464-06.

(24) Cheng, K.; Sun, Y.; Yu, H.; Hu, Y.; He, Y.; Shen, Y. *Staphylococcus aureus* SOS response: Activation, impact, and drug targets. mLife 2024, 3 (3), 343–366. DOI: 10.1002/mlf2.12137.

(25) Maiques, E.; Ubeda, C.; Campoy, S.; Salvador, N.; Lasa, I.; Novick, R. P.; Barbe, J.; Penades, J. R. β-lactam antibiotics induce the SOS response and horizontal transfer of virulence factors in *Staphylococcus aureus*. J Bacteriol 2006, 188 (7), 2726–2729. DOI: 10.1128/JB.188.7.2726-2729.2006.

(26) Bezrukov, F.; Prados, J.; Renzoni, A.; Panasenko, O. O. MazF toxin causes alterations in Staphylococcus aureus transcriptome, translatome and proteome that underlie bacterial dormancy. Nucleic Acids Res 2021, 49 (4), 2085–2101. DOI: 10.1093/nar/gkaa1292.

(27) Do, T.; Schaefer, K.; Santiago, A. G.; Coe, K. A.; Fernandes, P. B.; Kahne, D.; Pinho, M. G.; Walker, S. *Staphylococcus aureus* cell growth and division are regulated by an amidase that trims peptides from uncrosslinked peptidoglycan. Nat Microbiol 2020, 5 (2), 291–303. DOI: 10.1038/s41564-019-0632-1.

(28) Page, J. E.; Skiba, M. A.; Do, T.; Kruse, A. C.; Walker, S. Metal cofactor stabilization by a partner protein is a widespread strategy employed for amidase activation. Proc Natl Acad Sci U S A 2022, 119 (26), e2201141119. DOI: 10.1073/pnas.2201141119.

(29) Schaefer, K.; Matano, L. M.; Qiao, Y.; Kahne, D.; Walker, S. *In vitro* reconstitution demonstrates the cell wall ligase activity of LCP proteins. Nat Chem Biol 2017, 13 (4), 396–401. DOI: 10.1038/nchembio.2302.

(30) Kawai, Y.; Marles-Wright, J.; Cleverley, R. M.; Emmins, R.; Ishikawa, S.; Kuwano, M.; Heinz, N.; Bui, N. K.; Hoyland, C. N.; Ogasawara, N.;, et al. A widespread family of bacterial cell wall assembly proteins. EMBO J 2011, 30 (24), 4931–4941. DOI: 10.1038/emboj.2011.358.

(31) Jorasch, P.; Warnecke, D. C.; Lindner, B.; Zahringer, U.; Heinz, E. Novel processive and nonprocessive glycosyltransferases from *Staphylococcus aureus* and Arabidopsis thaliana synthesize glycoglycerolipids, glycophospholipids, glycosphingolipids and glycosylsterols. Eur J Biochem 2000, 267 (12), 3770–3783. DOI: 10.1046/j.1432-1327.2000.01414.x.

(32) Kiriukhin, M. Y.; Debabov, D. V.; Shinabarger, D. L.; Neuhaus, F. C. Biosynthesis of the glycolipid anchor in lipoteichoic acid of Staphylococcus aureus RN4220: role of YpfP, the diglucosyldiacylglycerol synthase. J Bacteriol 2001, 183 (11), 3506–3514. DOI: 10.1128/JB.183.11.3506-3514.2001.

(33) Zhang, B.; Liu, X.; Lambert, E.; Mas, G.; Hiller, S.; Veening, J. W.; Perez, C. Structure of a proton-dependent lipid transporter involved in lipoteichoic acids biosynthesis. Nat Struct Mol Biol 2020, 27 (6), 561–569. DOI: 10.1038/s41594-020-0425-5.

(34) Muscato, J. D.; Morris, H. G.; Mychack, A.; Rajagopal, M.; Baidin, V.; Hesser, A. R.; Lee, W.; Inecik, K.; Wilson, L. J.; Kraml, C. M.;, et al. Rapid inhibitor discovery by exploiting synthetic lethality. J Am Chem Soc 2022, 144 (8), 3696–3705. DOI: 10.1021/jacs.1c12697.

(35) Campbell, J.; Singh, A. K.; Santa Maria, J. P., Jr.; Kim, Y.; Brown, S.; Swoboda, J. G.; Mylonakis, E.; Wilkinson, B. J.; Walker, S. Synthetic lethal compound combinations reveal a fundamental connection between wall teichoic acid and peptidoglycan biosyntheses in Staphylococcus aureus. ACS Chem Biol 2011, 6 (1), 106–116. DOI: 10.1021/cb100269f.

(36) Brown, K. Crystal structure of the bifunctional N-acetylglucosamine 1-phosphate uridyltransferase from *Escherichia coli*: a paradigm for the related pyrophosphorylase superfamily. The EMBO Journal 1999, 18 (15), 4096–4107. DOI: 10.1093/emboj/18.15.4096.

(37) Massa, S. M.; Sharma, A. D.; Siletti, C.; Tu, Z.; Godfrey, J. J.; Gutheil, W. G.; Huynh, T. N. c-di-AMP accumulation impairs muropeptide synthesis in *Listeria monocytogenes*. J Bacteriol 2020, 202 (24). DOI: 10.1128/JB.00307-20.

(38) Devaux, L.; Sleiman, D.; Mazzuoli, M. V.; Gominet, M.; Lanotte, P.; Trieu-Cuot, P.; Kaminski, P. A.; Firon, A. Cyclic di-AMP regulation of osmotic homeostasis is essential in Group B *Streptococcus*. PLoS Genet 2018, 14 (4), e1007342. DOI: 10.1371/journal.pgen.1007342.

(39) Oberkampf, M.; Hamiot, A.; Altamirano-Silva, P.; Belles-Sancho, P.; Tremblay, Y. D. N.; DiBenedetto, N.; Seifert, R.; Soutourina, O.; Bry, L.; Dupuy, B.; et al. c-di-AMP signaling is required for bile salt resistance, osmotolerance, and long-term host colonization by *Clostridioides difficile*. Sci Signal 2022, 15 (750), eabn8171. DOI: 10.1126/scisignal.abn8171.

(40) Zeden, M. S.; Schuster, C. F.; Bowman, L.; Zhong, Q.; Williams, H. D.; Grundling, A. Cyclic di-adenosine monophosphate (c-di-AMP) is required for osmotic regulation in Staphylococcus aureus but dispensable for viability in anaerobic conditions. J Biol Chem 2018, 293 (9), 3180–3200. DOI: 10.1074/jbc.M117.818716.

(41) Kruger, L.; Herzberg, C.; Rath, H.; Pedreira, T.; Ischebeck, T.; Poehlein, A.; Gundlach, J.; Daniel, R.; Volker, U.; Mader, U.;, et al. Essentiality of c-di-AMP in *Bacillus subtilis*: Bypassing mutations converge in potassium and glutamate homeostasis. PLoS Genet 2021, 17 (1), e1009092. DOI: 10.1371/journal.pgen.1009092.

(42) Tosi, T.; Hoshiga, F.; Millership, C.; Singh, R.; Eldrid, C.; Patin, D.; Mengin-Lecreulx, D.; Thalassinos, K.; Freemont, P.; Grundling, A. Inhibition of the *Staphylococcus aureus* c-di-AMP cyclase DacA by direct interaction with the phosphoglucosamine mutase GlmM. PLoS Pathog 2019, 15 (1), e1007537. DOI: 10.1371/journal.ppat.1007537.

(43) Dengler, V.; McCallum, N.; Kiefer, P.; Christen, P.; Patrignani, A.; Vorholt, J. A.; Berger-Bachi, B.; Senn, M. M. Mutation in the C-di-AMP cyclase dacA affects fitness and resistance of methicillin resistant Staphylococcus aureus. PLoS One 2013, 8 (8), e73512. DOI: 10.1371/journal.pone.0073512.

(44) Dengler Haunreiter, V.; Tarnutzer, A.; Bar, J.; von Matt, M.; Hertegonne, S.; Andreoni, F.; Vulin, C.; Kunzi, L.; Menzi, C.; Kiefer, P.; et al. c-di-AMP levels modulate *Staphylococcus aureus* cell wall thickness, response to oxidative stress, and antibiotic resistance and tolerance. Microbiol Spectr 2023, 11 (6), e0278823. DOI: 10.1128/spectrum.02788-23.

(45) Lai, L. Y.; Satishkumar, N.; Cardozo, S.; Hemmadi, V.; Marques, L. B.; Huang, L.; Filipe, S. R.; Pinho, M. G.; Chambers, H. F.; Chatterjee, S. S. Altered PBP4 and GdpP functions synergistically mediate MRSA-like high-level, broad-spectrum beta-lactam resistance in *Staphylococcus aureus*. mBio 2024, 15 (5), e0288923. DOI: 10.1128/mbio.02889-23.

(46) Luo, Y.; Helmann, J. D. Analysis of the role of *Bacillus subtilis* sigma(M) in beta-lactam resistance reveals an essential role for c-di-AMP in peptidoglycan homeostasis. Mol Microbiol 2012, 83 (3), 623–639. DOI: 10.1111/j.1365-2958.2011.07953.x.

(47) Mukherjee, A.; Huang, Y.; Oh, S.; Sanchez, C.; Chang, Y. F.; Liu, X.; Bradshaw, G. A.; Benites, N. C.; Paulsson, J.; Kirschner, M. W.; et al. Bacterial cell wall biosynthesis is controlled by growth rate dependent modulation of turgor pressure in *E. coli*. bioRxiv 2025. DOI: 10.1101/2023.08.31.555748.

(48) Proseus, T. E.; Zhu, G. L.; Boyer, J. S. Turgor, temperature and the growth of plant cells: using Chara corallina as a model system. J Exp Bot 2000, 51 (350), 1481–1494. DOI: 10.1093/jexbot/51.350.1481.

(49) Sommer, A.; Fuchs, S.; Layer, F.; Schaudinn, C.; Weber, R. E.; Richard, H.; Erdmann, M. B.; Laue, M.; Schuster, C. F.; Werner, G.;, et al. Mutations in the gdpP gene are a clinically relevant mechanism for beta-lactam resistance in meticillin-resistant Staphylococcus aureus lacking mec determinants. Microb Genom 2021, 7 (9). DOI: 10.1099/mgen.0.000623.

(50) Witte, G.; Hartung, S.; Buttner, K.; Hopfner, K. P. Structural biochemistry of a bacterial checkpoint protein reveals diadenylate cyclase activity regulated by DNA recombination intermediates. Mol Cell 2008, 30 (2), 167–178. DOI: 10.1016/j.molcel.2008.02.020.

(51) Gandara, C.; Alonso, J. C. DisA and c-di-AMP act at the intersection between DNA-damage response and stress homeostasis in exponentially growing Bacillus subtilis cells. DNA Repair (Amst) 2015, 27, 1–8. DOI: 10.1016/j.dnarep.2014.12.007.

(52) Oppenheimer-Shaanan, Y.; Wexselblatt, E.; Katzhendler, J.; Yavin, E.; Ben-Yehuda, S. c-di-AMP reports DNA integrity during sporulation in *Bacillus subtilis*. EMBO Rep 2011, 12 (6), 594–601. DOI: 10.1038/embor.2011.77.

(53) Livak, K. J.; Schmittgen, T. D. Analysis of relative gene expression data using real-time quantitative PCR and the 2_-ΔΔCT_ Method. Methods 2001, 25 (4), 402–408. DOI: 10.1006/meth.2001.1262.

(54) Peneff, C.; Ferrari, P.; Charrier, V.; Taburet, Y.; Monnier, C.; Zamboni, V.; Winter, J.; Harnois, M.; Fassy, F.; Bourne, Y. Crystal structures of two human pyrophosphorylase isoforms in complexes with UDPGlc(Gal)NAc: role of the alternatively spliced insert in the enzyme oligomeric assembly and active site architecture. EMBO J 2001, 20 (22), 6191–6202. DOI: 10.1093/emboj/20.22.6191.

(55) Hopf, T. A.; Green, A. G.; Schubert, B.; Mersmann, S.; Scharfe, C. P. I.; Ingraham, J. B.; Toth-Petroczy, A.; Brock, K.; Riesselman, A. J.; Palmedo, P.;, et al. The EVcouplings Python framework for coevolutionary sequence analysis. Bioinformatics 2019, 35 (9), 1582–1584. DOI: 10.1093/bioinformatics/bty862.

(56) Crooks, G. E.; Hon, G.; Chandonia, J. M.; Brenner, S. E. WebLogo: a sequence logo generator. Genome Res 2004, 14 (6), 1188–1190. DOI: 10.1101/gr.849004.

(57) Schultz, B. J.; Snow, E. D.; Walker, S. Mechanism of D-alanine transfer to teichoic acids shows how bacteria acylate cell envelope polymers. Nat Microbiol 2023, 8 (7), 1318–1329. DOI: 10.1038/s41564-023-01411-0.

(58) Mirdita, M.; Schutze, K.; Moriwaki, Y.; Heo, L.; Ovchinnikov, S.; Steinegger, M. ColabFold: making protein folding accessible to all. Nat Methods 2022, 19 (6), 679–682. DOI: 10.1038/s41592-022-01488-1.

(59) Meng, E. C.; Goddard, T. D.; Pettersen, E. F.; Couch, G. S.; Pearson, Z. J.; Morris, J. H.; Ferrin, T. E. UCSF ChimeraX: Tools for structure building and analysis. Protein Sci 2023, 32 (11), e4792. DOI: 10.1002/pro.4792.

(60) Ducret, A.; Quardokus, E. M.; Brun, Y. V. MicrobeJ, a tool for high throughput bacterial cell detection and quantitative analysis. Nat Microbiol 2016, 1 (7), 16077. DOI: 10.1038/nmicrobiol.2016.77.

(61) Schindelin, J.; Arganda-Carreras, I.; Frise, E.; Kaynig, V.; Longair, M.; Pietzsch, T.; Preibisch, S.; Rueden, C.; Saalfeld, S.; Schmid, B.;, et al. Fiji: an open-source platform for biological-image analysis. Nat Methods 2012, 9 (7), 676–682. DOI: 10.1038/nmeth.2019.

(62) Kenny, M.; Schoen, I. Violin SuperPlots: visualizing replicate heterogeneity in large data sets. Mol Biol Cell 2021, 32 (15), 1333–1334. DOI: 10.1091/mbc.E21-03-0130.

(63) Schoch, C. L.; Ciufo, S.; Domrachev, M.; Hotton, C. L.; Kannan, S.; Khovanskaya, R.; Leipe, D.; McVeigh, R.; O’Neill, K.; Robbertse, B.;, et al. NCBI Taxonomy: a comprehensive update on curation, resources and tools. Database (Oxford) 2020, 2020. DOI: 10.1093/database/baaa062.

(64) Edgar, R. C. MUSCLE: multiple sequence alignment with high accuracy and high throughput. Nucleic Acids Res 2004, 32 (5), 1792–1797. DOI: 10.1093/nar/gkh340.

(65) Price, M. N.; Dehal, P. S.; Arkin, A. P. FastTree 2--approximately maximum-likelihood trees for large alignments. PLoS One 2010, 5 (3), e9490. DOI: 10.1371/journal.pone.0009490.

(66) Letunic, I.; Bork, P. Interactive Tree of Life (iTOL) v6: recent updates to the phylogenetic tree display and annotation tool. Nucleic Acids Res 2024, 52 (W1), W78–W82. DOI: 10.1093/nar/gkae268.

